# Nanopore device-based fingerprinting of RNA oligos and microRNAs enhanced with an Osmium tag

**DOI:** 10.1101/664169

**Authors:** Madiha Sultan, Anastassia Kanavarioti

## Abstract

Nanopores, both protein and solid-state, are explored as single molecule analytical tools, but using an experimental platform is challenging. Here we show that a commercially available nanopore device, MinION from Oxford Nanopore Technologies (ONT), successfully accomplishes a task challenging for a conventional analytical tool. Specifically the MinION discriminates among 31 nucleotide (nt) long oligoriboadenylates with a single pyrimidine (Py) substitution, when this pyrimidine is tagged/labeled with a bulky group (Osmium tetroxide 2,2’-bipyridine or OsBp). This platform also discriminates between an osmylated Py (Py-OsBp) followed by a purine (Pu) and a Py-OsBp followed by a second Py-OsBp, leading to the conjecture that the bulky tag enables sensing of a two-nucleotide sequence. Two-nucleotide sensing could greatly improve base-calling accuracy in motor enzyme-assisted nanopore sequencing.

We attribute the observed discrimination neither to the specific pore protein nor to OsBp, but to the tag’s bulkiness, that leads to markedly slower translocation and “touching” proximity at the pore’s constriction zone, that forces desolvation and reorganization, and enables strong interactions among the nanopore, the tagged pyrimidine, and the adjacent nucleobase. These results constitute proof-of-principle that size-suitable nanopores may be superior to traditional analytical tools, for the characterization of RNA oligos and microRNAs enhanced by selective labelling.

## Introduction

Nanopores made critical strides the last 30 years primarily due to the efforts of scientists and technologists who spearheaded nanopore-based DNA sequencing.^1-10^ In this approach a protein nanopore, with sub 2nm diameter, is inserted within an isolated membrane that separates two compartments filled with electrolyte. Applying a voltage across the two compartments leads to a constant flow of electrolyte ions via the nanopore (*i-t*), and this flow is partially blocked by the occasional passage of a single nucleic acid strand through the pore. Numerous studies using wt or bioengineered alpha-Hemolysin (α-HL)^11-16^ (Figure 1A), or a mutant version of the nanopore formed by Mycobacterium smegmatis porin A (MspA)^17,18^ illustrate that *i-t* conductance measurements (Figure 1B) yield ion current modulation with information attributed to nanopore/nucleobase interactions at the constriction zone of the pore (Figure 1C). Even though this technology avoids synthesis of the complementary strand, and hence enables direct sequencing, it is assisted by a processing enzyme that slows down the otherwise too fast translocation and acts as a motor to move the DNA/RNA strand one nucleobase at a time.^19-21^ Engineered versions of *α*–HL and MsPA with enhanced sensing capabilities report *i-t* data that vary as a function of a subsequence of, at least, 4 bases, and not as a function of a single nucleotide traversing the pore’s constriction point.^5^ Assuming that each 4-base subsequence exhibits a distinct ion current level, *I*_*r*_ (Figure 1C), there are 4^4^=256 *I*_*r*_ levels to be discriminated within approximately 80 picoAmperes (pA) of total measurement space^22^ with a standard deviation around ±2 pA, making base calling a challenging task. This issue has been addressed by utilizing sophisticated algorithms and matching the obtained sequence to a reference.^23,24^ The challenge is greater for RNA, compared to DNA, because RNAs, both coding and non-coding, are known to incorporate, in addition to the 4 canonical, some from a pool of currently known 150 post-transcriptionally modified nucleobases that fine tune RNA stability and function.^25^

**Figure 1.**
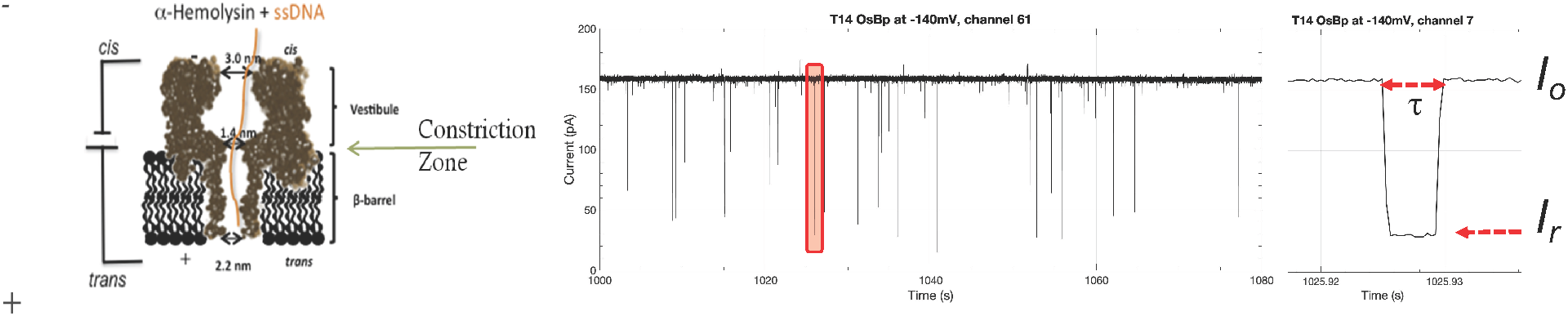
**Left**, Depiction of the wt *α*-Hemolysin (*α*-HL) pore showing the 1.4 nm constriction zone and the rather long but confined β-barrel known to fit a sequence of 10 bases held responsible for the observed residual ion current. For translocations using *α*-HL see Figure S17 in the Supplementary Information. **Middle**, raw 80s long *i-t* MinION recording illustrating translocations of a synthetic 31nt RNA T14 (sequence: A_14_GC(OsBp)GA_14_) at 34°C and at −140mV using ONT proprietary nanopore/buffer, but not ONT protocol. **Right**, Isolated single molecule translocation (pink block in middle figure) to show open pore ion current, *I*_*o*_, residual ion current of the translocation, *I*_*r*_, and dwell time or duration, *τ*.

In addition to sequencing, several nanopore platforms are successfully employed for single molecule analysis experiments. Aerolysin was shown to discriminate among short DNA oligos of different length.^26^ Solid-state nanopores were used to fingerprint nucleic acid nanoparticles.^27^ An engineered Fragaceatoxin C nanopore was shown to analyze a wide range of peptide lengths.^28^ Cytolysin A was shown to monitor the function of proteins when they are lodged inside the nanopore and report the concentration of glucose and asparagine directly from samples of blood, sweat, and other bodily fluids.^29^ α-HL was used to measure the kinetics of enzymatic phosphodiester bond formation.^30^ Using an engineered version of MspA a tool was developed to measure the movement of enzymes along DNA in real time.^31^ Noteworthy using an experimental platform is a complex process. Specifically, the monomeric unit of the nanopore protein needs to be manufactured, purified, assembled as a pore and inserted in a premade stable lipid bilayer, followed by addition of the nucleic acid in a small reaction vessel without disrupting the formation of the nanopore; all this before experimentation can begin.^5^ To expedite our work, we exploited the MinION, a commercially available small device from ONT, that has revolutionized DNA and RNA sequencing by allowing a worker with no other infrastructure besides a laptop and internet, to conduct sequencing experiments in any environment. The MinION is affordably priced, equipped with 512 preassembled nanopores from a proprietary engineered version of the CsgG protein,^32^ records simultaneously up to 512 *i-t* traces, and is set-up to visualize raw *i-t* data in a MatLab file format. The convenience of this device led us to evaluate it as a simple analytical tool for RNA oligo analysis and compare it with the resolution of traditional analytical tools like high-performance liquid-chromatography (HPLC). Experiments were conducted using as electrolyte the buffer provided by ONT for sequencing experiments, but no library was prepared, and no motor enzyme was included. Hence all the experiments reported here are voltage-driven, unassisted, single molecule translocations.

Under typical conditions unassisted voltage-driven translocation of RNA via a nanopore is very fast and measures 5 to 10 µs per base.^5^ The MinION exhibits a 3.012 kHz sampling rate, equivalent to reporting 3 data points per 1 ms. This slow acquisition rate implies that RNAs, let us say 100nt or shorter, may go “silent”, i.e. will not be detected by this device. Recently it was also reported that RNAs shorter than 200nt were not accessible to sequencing by the MinION using the recommended ONT protocol, i.e. with library preparation and incorporation of the motor enzyme.^33^ Hence the MinION doesn’t appear suitable for short RNA characterization. Current microarray, sequencing (Ion Torrent or Illumina (small RNA-seq)), and PCR-based methods for “small RNA” identification and/or quantitation are costly, complex, and have high time and resource demands.^34,35^ In addition, the most popular approaches are based on sequencing by synthesis and misread any modified RNA base. Since several regulatory RNAs are in the range from 22nt to 300nt long, it is important to find ways to characterize them and establish the presence, identity, and abundance of any post-transcriptionally modified bases within.

In order to characterize RNA oligos via the MinION and identify any modified RNA bases, we proposed to leverage the physicochemical properties/differences of the bases and selectively tag one or more with a bulky label.^36^ Assuming such label has high specificity, high selectivity, and exhibits negligible side-reactions, the bulkiness alone is expected to slow down translocation to measurable, readable *I*_*r*_ levels, as well as achieve discrimination, based on steric hindrance considerations, between a base conjugated to a bulky label and a native base. Such alteration to nucleic acid structure should reduce *i-t* dependence from a 4-base subsequence to, perhaps, a single labeled base. Importantly the selectivity/reactivity of the label for one base over another could be exploited and enable base identification for some among the 150 post-transcriptionally modified bases. Initial results using OsBp as pyrimidine-specific label demonstrated that expectations were mostly met, and showcased the potential as well as the limitations of the labeling approach and of this specific label.^37,38^

The reactivity of OsBp, known to add to the C5-C6 double bond of the pyrimidines, was established in the 1970s (Figure 2A).^39,40^ Recent development work showed that OsBp addition to DNA, coined here osmylation, using Yenos’ proprietary protocol is a remarkably clean reaction yielding pyrimidine conjugates in practically 100% yield with no detectable reactivity towards purines, and no phosphodiester bond cleavage.^41,42^ The selectivity of osmylation for deoxythymidine (dT) over deoxycytidine (dC), is 30-fold, and leads to labeling of either mostly dT, or both dT+dC using low or high OsBp concentration, respectively.^41^ Extensive studies with short, specifically designed DNA oligos, for which all products could be identified analytically by HPLC or capillary electrophoresis (CE), led to the conclusion that OsBp labeling is independent of sequence, length, and composition.^41^ Most importantly reactivity is not altered within a long sequence of pyrimidines, as evidenced by practically complete osmylation of dT_15_.^41^ A training set of oligos was exploited in developing a quantitative UV-vis assay that yields extent of pyrimidine osmylation, at a ±3% standard deviation, as a function of the ratio of pyrimidines over the total number of nucleotides in a strand (see later). As shown with M13mp18, a circular 7249nt ssDNA, secondary structure does not affect the efficiency of osmylation, most likely due to a denaturing effect exhibited by the OsBp reagent. Hence the same protocol yields practically 100% pyrimidine osmylation in short and long ssDNA.^41^

**Figure 2.**
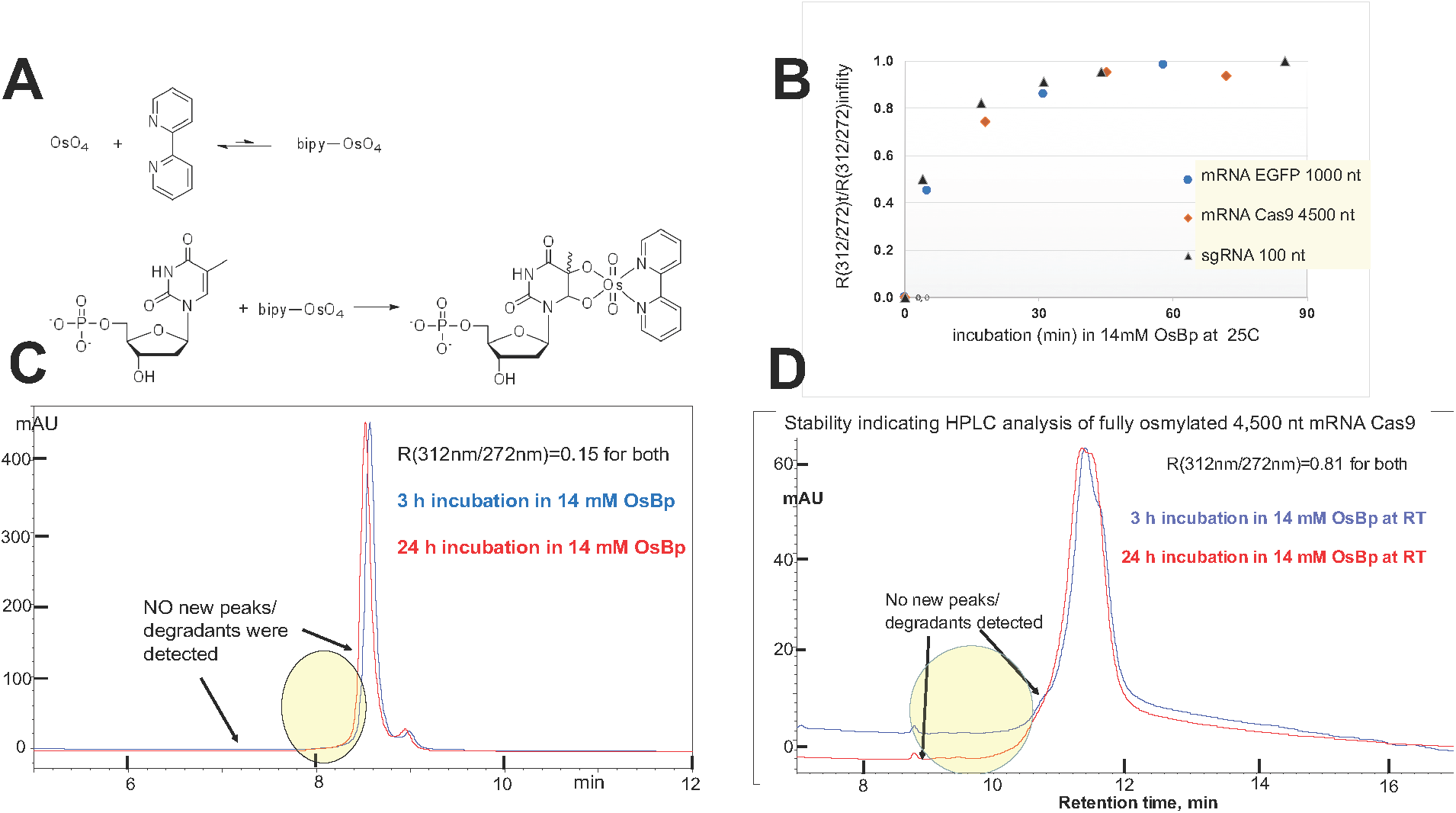
**A. Osmylation of Pyrimidines.** Preequibrium of Osmium tetroxide (OsO_4_) with 2,2’-bipyridine (bipy) to form a weak complex bipy-OsO_4_ or OsBp. In the following step OsBp adds to the C5-C6 double bond of a pyrimidine, thymidine monophosphate (TMP) shown here. Reaction of OsBp occurs from either side of the double bond and leads to topoisomers, often observed at 1:1 ratio. Due to the top/bottom addition the direction of the conjugate becomes parallel to the strand direction and the bulky OsBp extends all the way to the neighbor base. Evidence clearly shows that adjacent pyrimidines are kinetically as easily labeled as a monomer, as shown by dTTP being labeled as fast and as complete as oligo dT_15_.^41^ Even though labeling is not slowed down within a long sequence of pyrimidines, translocation via nanopores is (see later). **B. RNA Osmylation Kinetics.** Comparable kinetics (half-life of about 25 min) with 14 mM OsBp at 25° C were observed with a 100nt single guide RNA (sgRNA), a 1000nt mRNA EGFP and a 4500nt mRNA Cas9. Because the fraction of pyrimidines/ # of total nt is not equal for these three RNAs, observed absorbance ratio (R(312)/(272))_t_ at time t was normalized and plotted as a function of incubation time t. Normalization was done against the observed infinity value (R(312)/(272))_infinity_. **C. Stability-indicating HPLC profile at 260 nm of 74nt T6 (Table 1) in 14 mM OsBp for 3 or 24 h.** No new or increasing peaks were detected in the area in front of the main peak, consistent with undetectable degradation. Samples were quenched by removal of the excess label, and 3h samples were kept at −20°C until analysis, conducted at the same time as the 24 h samples. The T6 sample is rather concentrated, so that even degradants at 0.1% of T6 could be detected. IEX HPLC method at pH 12 (see Experimental Section). **D. Stability-indicating HPLC profile at 260nm of 4**,**500nt mRNA Cas9 in 14 mM OsBp for 3 or 24 h**, HPLC conditions as described in C. above.

The outstanding labeling features of OsBp led us to undertake nanopore-based single molecule translocation experiments in a number of collaborating Laboratories both in industry and in academia. Pore size suitability using solid-state silicon nitride (SiN) nanopores showed that 1.6nm wide SiN pores permit translocation of 80nt long osmylated deoxyoligos, and exhibit dramatic tranlocation slowdown with increasing osmylation.^38^ Personal communication from Professor Mark Akeson of the Genomic Institute of the University of California in Santa Cruz indicated that the same 80nt oligos translocate via the 1.4nm wide wt α-HL nanopore, albeit very slowly. Experiments with short, specifically designed oligodeoxynucleotides dA_10_XdA_9_ (X=deoxypyrimidine) via wt α-HL showed readably slow and distinct median translocation features for different X. Specifically when X = dA, dT(OsBp), dC(OsBp), dU(OsBp), or 5-MedC(OsBp) the distribution of the fractional residual ion current, *I*_*r*_*/I*_*o*_, where *I*_*o*_ is open pore ion current and *I*_*r*_ is the residual ion current of the blockade or event (Figure 1C), has a maximum at 0.14, 0.08, 0.11, 0.12 and 0.12 (with STD ±0.01), respectively, while observed duration or dwell times (τ) (Figure 1C) measured at −120mV are 50, 150, 310, 360 and 470µs, respectively.^37^ The observed effects are small with respect to residual ion current differences, but dramatically large with respect to translocation duration compared to 3 to 5µs per intact base,^5^ indicating a remarkable slowing down. These data suggest that the α-HL constriction zone interacts with both moieties, the OsBp and the nucleobase within a specific Py(OsBp), and illustrate proof-of-concept for single osmylated deoxypyrimidine differentiation in *α–*HL. The above studies clearly demonstrated that size-suitable nanopores, solid-state or protein, discriminate between a native and a single osmylated deoxypyrimidine during unassisted voltage-driven oligo translocation.

**Table 1:**
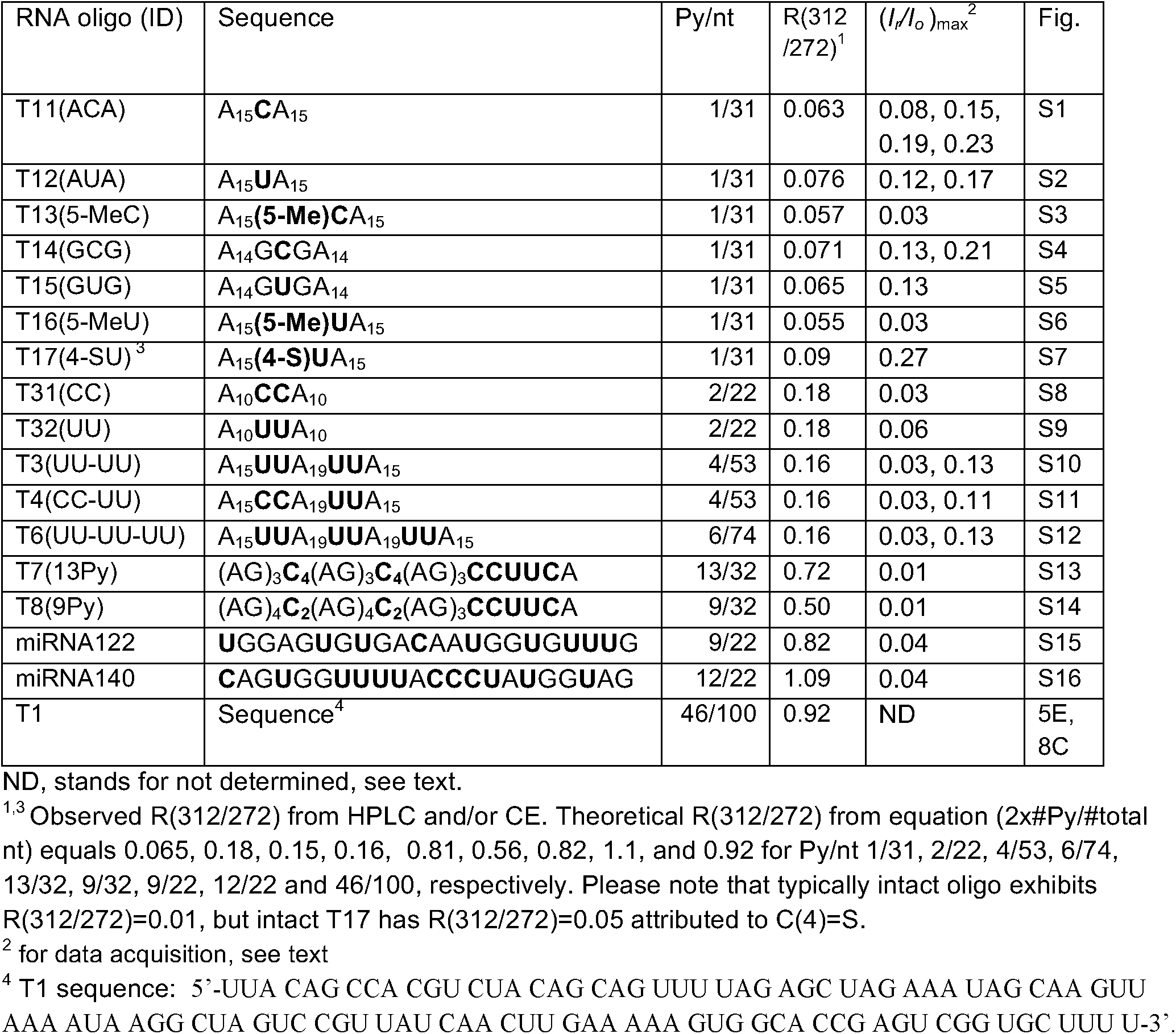
List of tested RNA oligos. RNA oligo ID, sequence (all ribonucleotides), number of pyrimidines over total nucleotides (Py/nt), observed Absorbance ratio (R(312/272)) of the osmylated derivative (see text), maxima of fractional residual ion current (*I*_*r*_*/I*_*o*_)_max_ and corresponding figure in the Supplementary Information.

RNA oligos in the range of 20 to 300nt include coding and non-coding nucleic acids with known functions in mRNA translation and in regulation of most biological processes.^43^ microRNA (miRNA) is a group of short RNAs, about 22nt long, that serves as a universal regulator with applications in personalized medicine as biomarker and as potential therapeutic.^44-48^ Due to the interest in short RNA characterization we extend here osmylation to RNA, and report that OsBp exhibits comparably excellent labeling properties with RNA pyrimidines, as described above for DNA pyrimidines. Using high quality synthetic RNA oligos (see Table 1) and the commercially available MinION we show that unlabeled 31nt oligos are not detected by this device, but that the corresponding osmylated oligos translocate unassisted via the CsgG proprietary nanopore protein of this device and are easily resolved from the noise and the instrument-generated lines, even when the oligo contains a single OsBp moiety. The reported fractional residual level of ion current, *I*_*r*_*/I*_*o*_, illustrate that each RNA is fingerprinted via the MinION and this fingerprint is visually distinct for 31nt RNAs with MW of 10,500 that differ by a single methyl group, a distinction hard to achieve by any other means. Close inspection of these *I*_*r*_*/I*_*o*_ fingerprints as a function of structural differences between the tested RNAs leads to the conjecture that the observed discrimination is not a feature of the specific protein pore or of the specific label used here, but it is the result of the bulkiness of a tag, that leads to “touching” proximity and markedly slower translocation at the pore’s constriction zone, that enables desolvation and reorganization as well as strong interactions between the nanopore, the tagged pyrimidine, and the adjacent nucleobase. Despite the stochastic nature of the translocation process, these interactions yield rudimentary information in the form of Py-Pu sequence, that may be sufficient for short RNA characterization in a mixture. Our data serve as proof-of-principle for exploring nanopores as analytical tools for short RNAs and miRNAs.

## RESULTS and DISCUSSION

### Labeling RNA pyrimidines with an Osmium tag: Manufacturing and Stability

Osmylation conditions for DNA were developed earlier;^40^ these conditions were reevaluated, and optimized for RNAs. Briefly the RNA osmylation protocol requires 3 hours incubation at room temperature in water in the absence of buffer in glass vials in the presence of 12 to 14mM OsBp (1:1 equimolar mixture of OsO_4_ and 2,2’-bipyridine) and yields practically 100% osmylation of the pyrimidines, even with long RNAs, like mRNA Cas9, known to exhibit secondary structure (Figure 2B). It is important that the ratio of mM OsBp to mM pyrimidine, in monomer equivalents, is 30-fold or more, so that the labeling is not slowed down by depletion of OsBp, while it reacts with the oligo or evaporates (see Experimental Section). The high excess of OsBp reagent to pyrimidine monomer is also required due to the low association constant between OsO_4_ and 2,2’-bipyridine as shown by square dependence of the osmylation kinetics on the “OsBp” concentration (Table S1 and Figure S17 in the Supplementary Information).

Osmylation kinetics at the above recommended conditions are independent of RNA length, composition, and secondary structure, as seen in Figure 2B. Specifically three RNAs of vastly different length (a 100nt long single guide RNA (sgRNA), a 1000nt long RNA, mRNA EGFP, and a 4500nt long RNA, mRNA Cas9) conform onto the same kinetics with about 25 min half-life. The observation, that even nucleic acids with secondary structure, such as mRNAs, label kinetically equally fast and as efficiently as an oligo, is likely due to a denaturing effect of OsBp solutions at concentrations as high or higher than 12mM.^41^ It turns out that in dilute, let us say, 5mM OsBp solutions only regions exposed to the solvent are labeled, while hybridized regions remain intact. Manufacturing conditions exploit high OsBp reagent concentration (12 to 14mM), and about 6 half-lives, so that the osmylation is practically 100%, and process duration is minimized. On the contrary, in order to determine selectivity, the osmylation kinetics are conducted at a lower concentration of OsBp (3 to 6mM). Osmylation kinetics were monitored automatically every 15 min using CE, and the CE profiles are included in the corresponding figure of the oligo in the Supplementary Information. With the exception of the oligoriboadenylate with 5-MeU substitution, which is subject to very fast osmylation, just as with the deoxy derivative,^41^ all other tested oligos exhibit comparable reactivity. Observed relative reactivity towards osmylation obtained from the kinetics of T11 though T16 using 5.2mM OsBp at 26°C in water are U/C=4.7, 5-MeC/U=0.9, 5-MeC/C=4.1 and 5-MeU/C=44, in excellent agreement with the relative reactivity observed in deoxy oligos.^37^ Osmylation reactivity depends on the electrophility of the C5-C6 double bond, and this is the feature that will enable selectivity of one vs. another nucleobase among some of the post-transcriptionally modified RNA bases. Typically the reactivity of the base in a sequence mirrors the reactivity of the mononucleotide.

Removal of the excess OsBp after manufacturing takes about 7 minutes, and is conducted using a TrimGen mini-column following the manufacturer’s instructions. Extent of purification can be assessed by CE or HPLC (for methods see Experimental Section), because OsBp migrates (CE) or elutes (HPLC) well ahead of the oligo and of the osmylated product. HPLC or CE profiles for each tested intact oligo and each fully osmylated oligo are included in the corresponding figure in the Supplementary Information; osmylation does not lead to side-reactions, as can be qualitatively seen in these profiles. In addition, the stability of the RNA in the presence of OsBp was evaluated with an extra pure 74nt long RNA and with a 4500nt long mRNA Cas9 using a stability-indicating HPLC method^49^ (see Experimental Section). Comparison between the HPLC profiles after 3 h and 24 h of osmylation (see Figure 2C with 74nt long RNA and Figure 2D with mRNA Cas9) illustrates no detectable changes, suggesting no detectable degradation during an additional 21 h prolonged incubation under manufacturing conditions.

Selective labeling of a nucleic acid requires an assay for quality control. It turns out that addition of OsBp to the C5-C6 Py double bond and formation of Py(OsBp) creates a new chromophore in the wavelength range of 300 to 320nm, where nucleic acids exhibit negligible absorbance. We exploited this observation and used a deoxyoligo training set to show that extent of osmylation can be measured using the equation R(312/272) = 2×(# of osmylated pyrimidines/total # of nucleotides), where R(312/272) is the ratio of the observed absorbance at 312nm over the observed absorbance at 272nm.^41^ Using the ratio R instead of the absorbance at 312nm serves to normalize the measurement, and minimize sampling errors. When experimental R(312/272) = 2× (# of pyrimidines/total # of nucleotides), osmylation is practically 100% complete.^41^ The wavelengths 312 nm and 272 nm were chosen in order to maximize the effect and to equalize contributions by osmylated dT, dC or dU, assuring that nucleic acid composition doesn’t affect the assay’s accuracy which stands at ± 3%. The R(312/272) data obtained in this study (Table 1) confirm that the above equation is also valid for osmylated RNA with bases U, C, 5-MeU, and 5-MeC. Notwithstanding the UV-vis assay should be tested and confirmed or modified for other non-canonical bases. Using HPLC and/or CE analysis the label/OsBp peak appears very early and resolves well from the osmylated oligo. However Nanodrop measurements will not differentiate between label and oligo and therefore removal of the label, which absorbs at 312nm, ahead of the measurement is necessary.

### HPLC analysis of RNA oligos

HPLC analysis is routinely used for identification, resolution from impurities, stability evaluation, and quality control of pharmaceuticals. RNAs, in the form of single guide RNA (sgRNA), mRNA, or miRNA, are being considered for therapeutic applications, and validated analytical methods to characterize them are being sought.^49^ Typically RNA oligos up to the 60-mer can be resolved by HPLC; both ion exchange (IEX)^50^ and ion-pair reversed phase (IP-RP)^51^ chromatographies are applicable and exploited here for analysis of the RNAs used in this study. As HPLC analytical columns (see Experimental Section) we used the ones that in our experience give the best possible resolution, but alternative columns/methods may be put to the test. Among the tested articles the seven 31nt oligos vary minimally from each other, and their separation and identification by HPLC is challenging. Despite method development efforts complete separation was not achieved; Figure 3A (top HPLC profile) using an IP-RP method illustrates that 6 out of the 7 31nt oligos appear as a single peak and Figure 3A (bottom HPLC profile) using an IEX method illustrates that 4 out of the 7 31nt oligos do not achieve base line resolution; hence the search for more suitable analytical methods continues. It should be noted that osmylated RNAs exhibit less resolution by HPLC compared to intact, because the OsBp moiety yields peak broadening and therefore intact oligos that are resolvable may overlap after osmylation.

**Figure 3.**
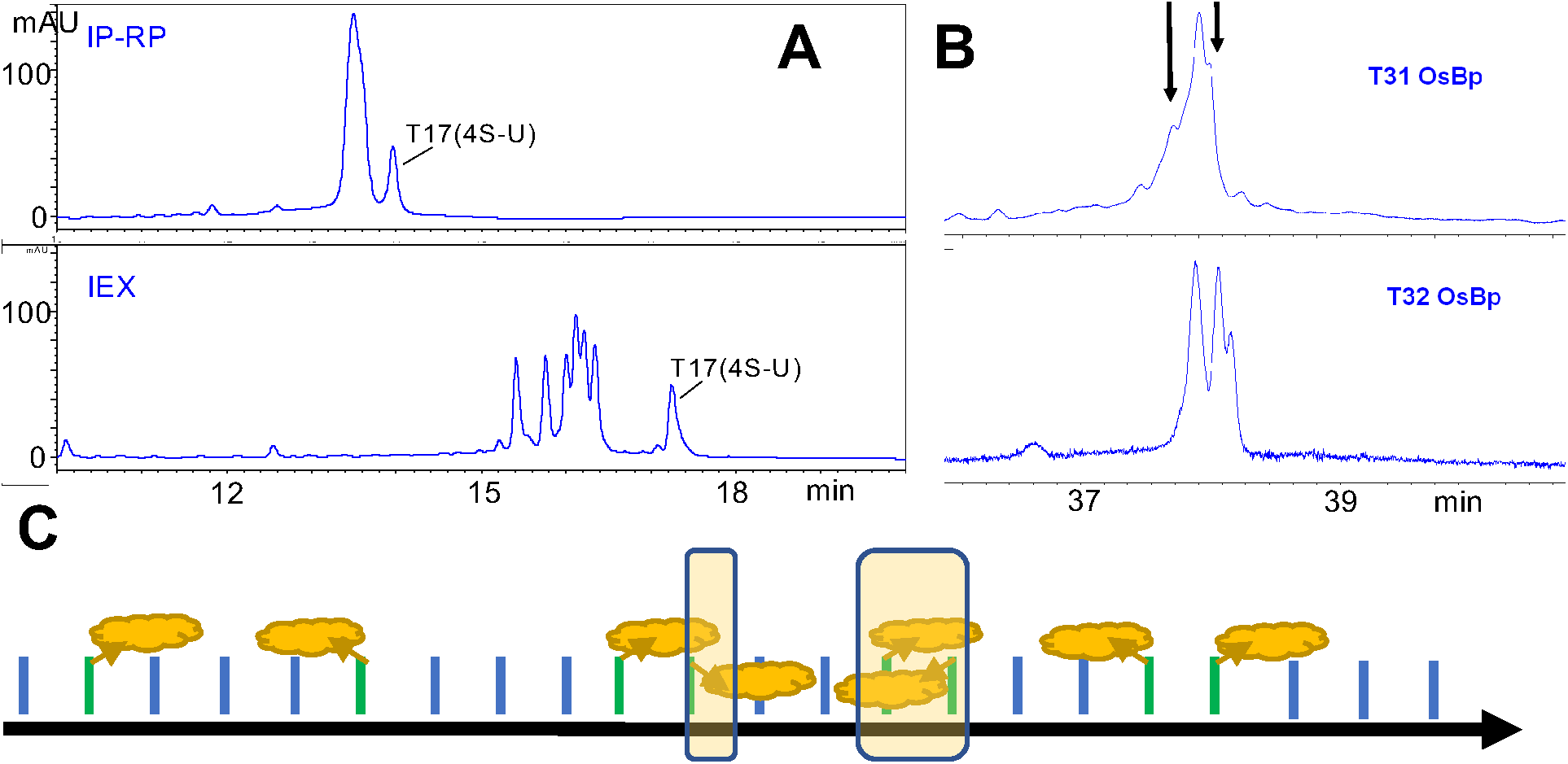
**A. HPLC profiles of an equimolar mixture of 31nt intact RNAs T11 through T17** (see Table 1) with minimal sequence differences, run by two methods, i.e. ion-pair reversed phase (IP-RP) (top) and ion-exchange (IEX) (bottom), to show that only T17(4-SU) resolves clearly and the other six RNAs elute closely together. **B. CE profiles of the osmylated T31(CC) and T32(UU) show the presence of three diastereomers.** The arrows in T31 indicate that the shoulders around the main peak are considered to be the two isomers. For statistical reasons the theoretical ratio is PA:OL:AP = 2:1:1 (see cartoon in C here) and observed ratio is close to the theoretical, only that PA configuration appears to migrate in the middle with T31 and first with T32. **C. Cartoon to show an RNA strand with 5’3’ direction** (black arrow), the two isomers for a single Py(OsBp) in a strand (Py (green bars), Pu (blue bars), OsBp (yellow cloud)), and the three isomers for a sequence with two consecutive Py(OsBp); the shaded rectangle approximates the extent of overlapping OsBp moieties for the three configurations and illustrates that overlap exists for the PA configuration, overlap is extensive for the OL configuration, and unlikely for the AP configuration. Depending on the relative direction of the PA configuration in the context of strand direction this configuration is 2-times more abundant compared to the other two. Please note that within the <u>least crowded</u> arrangement there is one doublet in the PA, two doublets in the PA, and one doublet in the OL configuration, with three, four, or five adjacent Py, respectively. This implies that RNAs that carry five adjacent Py at the 3’end should exhibit comparable translocation features as T7 and T8 (Figure 6H).

### OsBp’s topoisomerism

OsBp addition to the pyrimidine C5-C6 double bond occurs from either the top or from the bottom of the pyrimidine ring yielding two topoisomers.^40,41,52^ Due to the strand directionality these two isomers are detectable by either HPLC or CE (see profiles of osmylated 31nt oligos in Supplementary Information, part C). Two peaks were observed for every singly osmylated oligo tested here with a product ratio not far from unity, so that both are detectable. Since voltage-driven translocation of nucleic acids via nanopores is known to yield different *I*_*r*_*/I*_*o*_ levels depending on the entry direction of the strand, 3’ entry *vs* 5’ entry,^53^ it is plausible that a nanopore may exhibit up to four different *I*_*r*_*/I*_*o*_ levels for the four ways that a singly osmylated oligo can interact with the pore. In the absence of experiments with immobilized RNAs within the pore^54,55^ it is difficult to assess this proposition. Nevertheless the *I*_*r*_*/I*_*o*_ histogram of T11(AC*A) presents four maxima, (*I*_*r*_*/I*_*o*_)_max_, supporting the postulate of four *I*_*r*_*/I*_*o*_ levels in this case (FigS1, Supplementary Information).

OsBp’ isomerism and unhindered reactivity towards consecutive bases leads to three diastereomers in an oligo with two adjacent osmylated pyrimidines, as evidenced by CE in the form of 3 product peaks using osmylated T31(CC) and T32(UU) (see Table 1 and Figure 3B). Figure 3C is a cartoon to illustrate topoisomers from a single or from two adjacent pyrimidines. With two adjacent pyrimidines diastereomers result from (i) overlapping OsBp moieties (OL), i.e. lining antiparallel and towards each other, (ii) OsBp moieties lining parallel to each other (PA), and (iii) OsBp moieties lining antiparallel and away from each other (AP). Statistically the distribution of OL:PA:AP is 1:2:1 and the observations with osmylated T31(CC) and T32(UU) support it (Figure 3B), even though the order of migration appears different for these two materials.

### Nanopore Experiments

Motivated by the single Py(OsBp) discrimination in deoxyoligos observed with *α*-HL, experiments were extended to RNAs, initially using the NanoPatch instrument from Electronic Biosciences (EBS) equipped with a proprietary glass nanopore membrane (GNM) (see Experimental Section). In this platform (NanoPatch/wt α-HL) the discrimination of the nanopore for RNA compared to DNA appeared to be much stronger (Figure S18 in the Supplementary Information). To speed up the work, we tested the MinION from ONT, observed translocations using the osmylated RNAs, and present here this work. The flow cell of the MinION is a consumable, equipped with a total of 512 premade nanopores from a proprietary engineered version of the CsGg protein. The sequences of the tested RNA oligos 22nt to 100nt long are listed in Table 1. Their properties, as tested by HPLC, CE, and the MinION, are also summarized in Table 1. HPLC profile of the intact oligo, CE profiles of the osmylation kinetics at low OsBp concentration, the HPLC or CE profile of the osmylated oligo, as well as the histogram of the *I*_*r*_*/I*_*o*_ data determined for each can be found in the corresponding figure in the Supplementary Information. It should be emphasized that the ONT protocol was not followed, no library was created, and the processing enzyme was not included in this study. Hence all the observed translocations are unassisted, driven simply by the voltage drop, typically conducted with a biased voltage in the range −140mV to −220mV.

Figure 4 represents concatenated *i-t* traces from three experiments each conducted with a 31nt RNA, an oligoribo-adenylate with a single Uridine derivative in the middle of the strand, i.e. 5-MeU, U, and 4-SU. Visual inspection, as highlighted by the colored boxes, suggests that the observed translocations with the oligo carrying 5-MeU(OsBp) yield substantially less residual ion current compared to the translocations with the oligo carrying U(OsBp), and the later yields dramatically less residual ion current compared to the one with the 4-SU(OsBp). The “*I*_*r*_” value at the middle of each box is around 20, 40, and 100pA for 5-MeU, U, and 4-SU, respectively, spanning a range of 80pA, somewhat larger than what currently is used for sequencing RNA with the MinION.^22^ These data support nanopore-based single Py(OsBp) discrimination in RNAs and this discrimination is made visible using the MinION platform.

**Figure 4.**
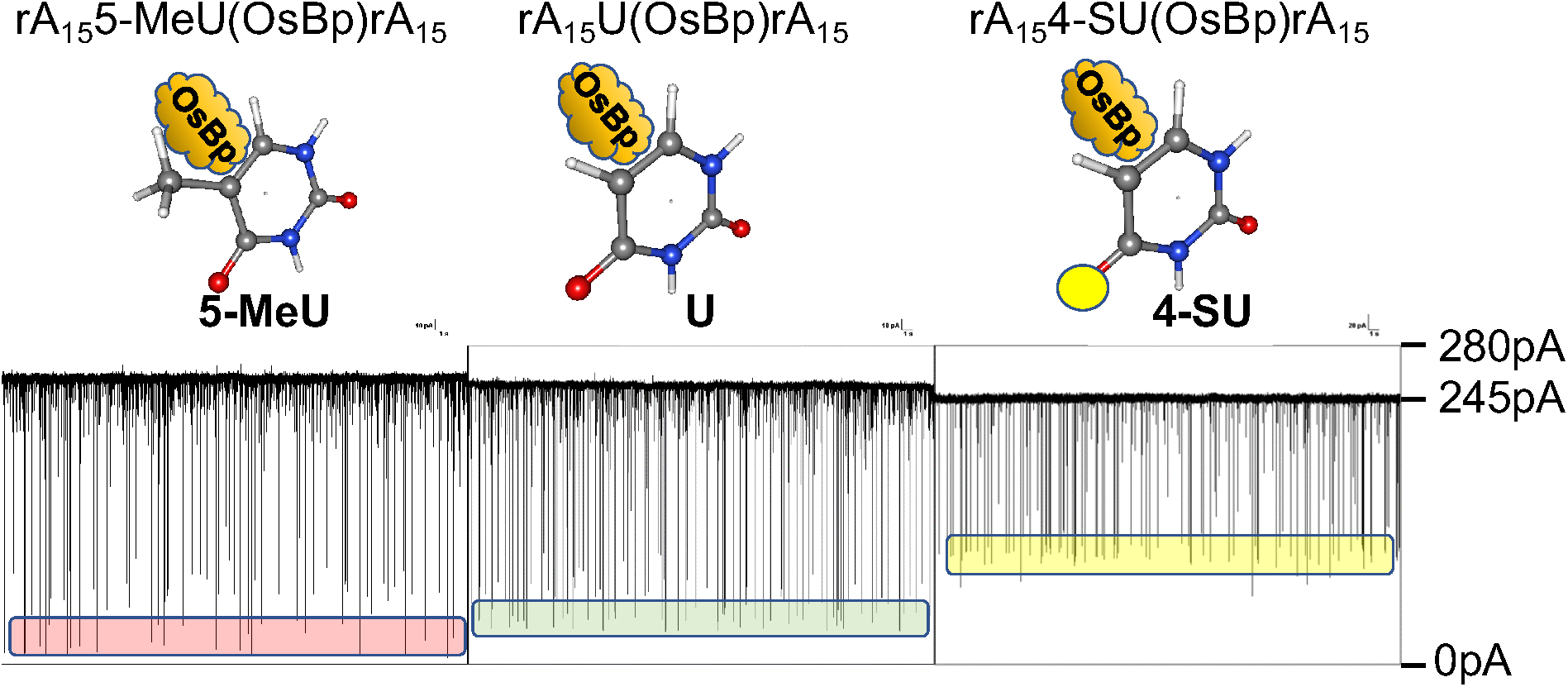
Visible discrimination in 31nt RNAs from concatenated *i-t* traces for unassisted voltage-driven single molecule translocations using the MinION device from ONT; each *i-t* trace spans about 100s. Please note that these three are the only concatenated MinION *i-t* traces presented in this report. From left to right, *i-t* traces shown for osmylated 31nt RNAs: T16(5-MeU), T12(U) and T17(4-SU), see top for sequences and below cartoon to show the osmylated pyrimidine base moiety with OsBp at the C5-C6 bond. The presence of a 5-Me moiety on the U of this RNA which is practically an oligoadenylate yields the most ion current obstruction (pink block), less ion current obstruction with U (green block), and remarkably less ion current obstruction with a 4-SU moiety (yellow block); see text for discussion and Table 1 for quantitation of the effect.

### Qualification of the MinION as an analytical tool

To the best of our knowledge, the MinION is exclusively used for enzyme-assisted sequencing. Hence our work regarding voltage-driven RNA characterization is a novel application for this platform. In this context, we evaluated MinION’s suitability as an analytical tool for short RNAs. Three basic questions were addressed: (a) Do all working nanopore channels provide comparable results, (b) Are intact RNA oligos detected by the MinION, and (c) Is the MinION’s pore protein a size-suitable nanopore for osmylated RNA. Briefly the answers to these questions are: (a) and (c) Yes; (b) No for shorter than 31nt and Yes for longer than 74nt RNA. The first question was answered by visually inspecting every recorded *i-t* trace from each working channel and by graphing the translocation events (see discussion below) separately for a number of channels. Judging from obtaining superimposable histograms from different channels, channels are comparable. Open pore ion current (*I*_*o*_) from different channels may vary by up to ±15 pA within a single experiment, but data are normalized by reporting the fractional residual ion current (*I*_*r*_*/I*_*o*_) as seen in the histograms in the Supplementary Information. Typically two experiments were conducted with the same oligo using different flow cells, and data are reported from 4 or more channels for each experiment. It was practically established that 300 *I*_*r*_*/I*_*o*_ values are sufficient to yield reproducible results, and 500 to 1400 *I*_*r*_*/I*_*o*_ values are reported per oligo.

The question whether or not the MinION reports translocations from intact RNA was evaluated by experiments conducted in the absence of any oligo, and in the presence of intact RNAs. Figure 5A shows the raw *i-t* recording of ONT proprietary buffer. It is noticeable that the open pore current is interrupted by “instrument lines” only, and not by any translocation events. These “instrument lines” traverse the *i-t* trace, and typically exhibit a mirrored image of lines extending vertically up and down. Figure 5B illustrates the raw *i-t* recording from an experiment conducted with an intact 31nt RNA (T11(ACA)) that appears identical to the one in Figure 5A. The experiment was conducted at the typical concentration and at a 3-fold higher concentration (shown in Figure 5B) and no translocations or other detectable difference were observed confirming that this oligo is not detected by the pore. The observation suggests that 31nt RNAs or shorter translocate faster than the instrument’s ability to record them, and it is in agreement with the instrument’s specifications of 3 data points per 1 ms, as discussed earlier. In contrast, Figure 5C presents the raw *i-t* recording from an intact 74nt RNA and illustrates the presence of a large number of translocations, accompanied by a small number of “instrument lines” (not shown here due to the choice of y-axis). Some of the events exhibit high residual current *I*_*r*_, and some exhibit low *I*_*r*_. The former are many and considered to be bumping events of the RNA on the pore; the later are fewer and considered to be actual RNA translocations. Within the range of the true translocations events, i.e. below 50pA in Figure 5C, one visually identifies the existence of two *I*_*r*_ levels, as shown by the two blue transparent blocks. We attribute these two *I*_*r*_ levels by extrapolation to the reported distinct translocation features by a 3’-entry or a 5’-entry of an oligo via a nanopore.^53^ Figure 5C clearly suggests that intact 74nt RNAs or longer are easily detected by the MinION. We didn’t probe this issue any further, as recording of intact oligos in the range between 31nt and 74nt may depend on sequence. Notably intact poly(C) and poly(U) translocate via the MinION and exhibit substantially lower *I*_*r*_ levels (*I*_*r*_*/I*_*o*_ ≈ 0.05, 0.10), compared to the one observed with the 74nt RNA (*I*_*r*_*/I*_*o*_ = 0.15, see below and in Figure 6E) which is practically an adenylate, in agreement with early reports.^1,2^

**Figure 5:**
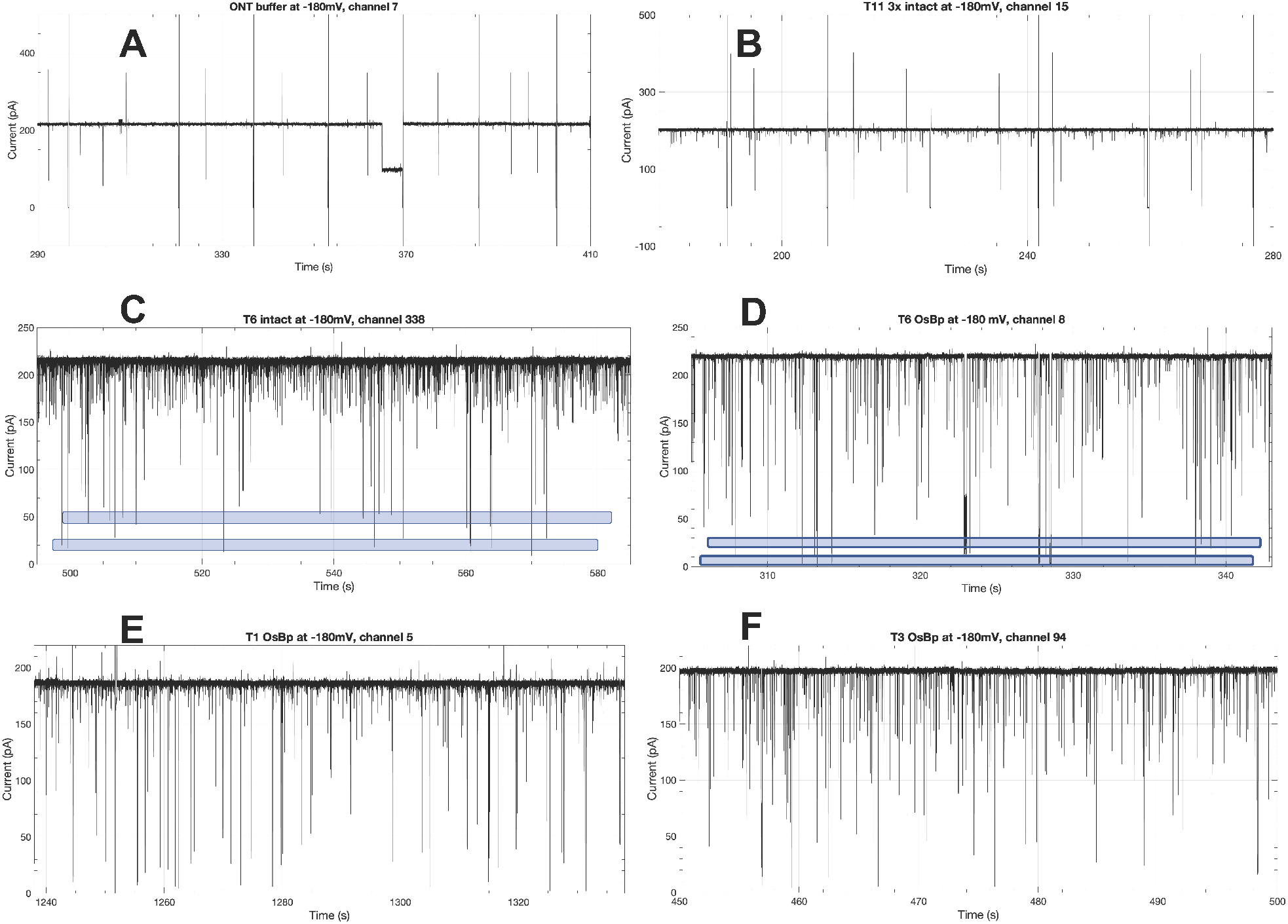
Visual discrimination of osmylated RNA oligos by the MinION. Raw *i-t* MinION recordings from different experiments conducted at 34°C (not a variable) and at −180mV. **A.** ONT buffer alone, 120 second trace, observed lines are device or instrument-generated and appear more often in the absence of any translocating molecules. **B.** Intact 31nt RNA T11 at 3× the typical concentration, 100 second trace, appears identical to A indicating that T11 translocations are likely too fast and are not recorded by the MinION. **C.** Intact 74nt RNA T6, 90 second trace, showing a lot of events with high residual current, attributed to bumping events (see text) and a number of events with low residual current ion (two blue blocks). The latter suggests that translocations of intact RNAs as long as 74nt are recorded by the MinION. **D.** Same RNA as in C but osmylated (6 OsBp), 40 second trace, showing that the osmylated oligo exhibits more current ion obstruction compared to the intact, as evidenced by the lower level of these blue blocks compared to the ones with the intact; quantitation in Table 1. **E.** Osmylated T1, a 100nt sgRNA with known secondary structure becomes linearized and traverses the pore, 50 second trace, indicating that RNAs as long as 100nt can be characterized using our osmylation/unassisted translocation approach via the MinION. **F.** Osmylated T3, a 53nt RNA, 50 second trace, at the same µM concentration as T1 (F*). i-t* trace of 100nt T1 exhibits remarkably fewer high residual ion current events (*I*_*r*_ > 125 pA) compared to *i-t* trace of 53nt T3 and 74nt T6, suggesting that oligoadenylates may artificially produce more bumping events.

**Figure 6.**
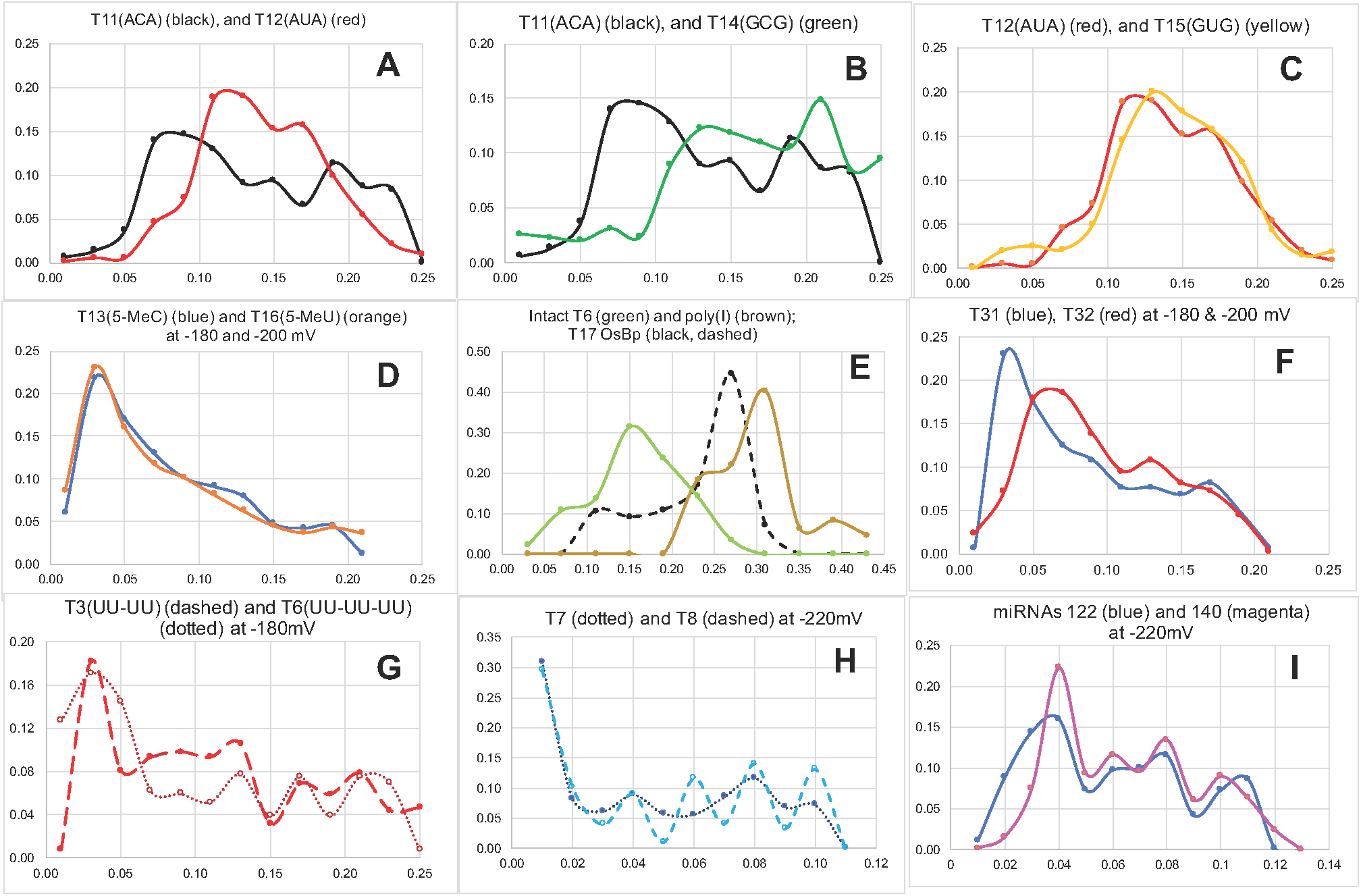
Plots of normalized *I*_*r*_*/I*_*o*_ Histograms from unassisted translocation of osmylated RNAs via the MinION nanopore device; x-axis, *I*_*r*_*/I*_*o*_ with bin size=0.01, 0.02 or 0.04; y-axis, normalized counts per bin by total counts reported. Osmylation of pyrimidines practically 100%. x,y axes vary to match the specific set of RNAs. Values (*I*_*r*_*/I*_*o*_)_max_ reported in Table 1, actual *I*_*r*_*/I*_*o*_ histograms in the corresponding figure in the Supplementary Information. For discussion see text. **A.** T11(ACA) vs. T12(AUA) at −180mV exhibit distinct profiles. **B.** T11(ACA) vs. T14(GCG) at −180mV exhibit distinct profiles. **C.** T12(AUA) vs. T15(GUG) at −180mV exhibit comparable profiles. **D.** T13(5-MeC) vs. T16(5-MeU) exhibit identical profiles. **E.** T17(4-SU) vs. 74nt intact T6 vs. poly(I) to show that (*I*_*r*_*/I*_*o*_)_max_ increases in the order T6 intact <osmylated T17<intact poly(I). **F.** T31(CC) vs. T32(UU) exhibit distinct profiles, and comparable to the trends observed in A, but shifted to relatively more ion current obstruction. **G.** 53nt T3(UU-UU) vs. 74nt T6(UU-UU-UU) exhibit comparable profiles, only that translocations with the longer oligo are shifted towards more obstruction compared to the translocations of the shorter oligo (see text). **H.** 32nt RNAs T7(13Py) vs. T8(9Py), of similar sequence but different number of Py(OsBp), exhibit comparable profiles, and show the most ion current obstruction seen in this study, attributed to a sequence of 5 adjacent osmylated pyrimidines. **I.** 22nt oligos, miRNA122(9/22) vs. miRNA140(12/22) exhibit comparable profiles for translocations at relatively low ion current obstruction, but distinct profile at the highest ion current obstruction, indicating substantially different subsequences of multiple adjacent pyrimidines (see discussion in text). Notably (*I*_*r*_*/I*_*o*_)_max_=0.04 for the two miRNAs is in between the (*I*_*r*_*/I*_*o*_)_max_=0.03 and 0.06 of the T31(CC) and T32(UU) respectively, in agreement with their sequences containing both C and U bases.

Figure 5D represents the raw *i-t* recording from a fully osmylated 74nt RNA with six U(OsBp) moieties showing the presence of a large number of events. Again here most events exhibit high residual current *I*_*r*_, attributed to “bumping events”, and some exhibit low *I*_*r*_, attributed to true RNA(OsBp) translocations grouped into two *I*_*r*_ levels, as seen by the blue transparent blocks. It is noteworthy that both *I*_*r*_ levels with the osmylated oligo are below the *I*_*r*_ levels of the intact, suggesting that a distinction can be visually made for experiments with intact vs. osmylated oligo, as observed in the experiments with deoxyoligos via *α*-HL, and attributed to the presence of the bulky OsBp tag within the confined space of a nanopore.

To answer the third question regarding suitability of the MinION’s pore for osmylated RNAs, we tested for oligo length and pyrimidine composition to assess (a) whether relatively long oligos, such as the 100nt sg RNA(OsBp) can traverse the pore, and (b) whether the presence of multiple consecutive Py(OsBp) prohibits translocation. Figure 5E shows a 50s raw *i-t* trace from an experiment conducted with a fully osmylated 100nt RNA (see sequence in Table 1). It is noteworthy that this 100nt RNA is shown to exhibit distinct secondary structure at elevated temperatures,^49^ and the intact molecule is not expected (not tested yet) to traverse the pore. As mentioned earlier osmylation linearizes the strand, most likely due to interruption of base-stacking, once the pyrimidines are osmylated, and this should enable translocation of long RNA, most likely of any length. In contrast to Figure 5E, Figure 5F shows a 50s long raw *i-t* trace from osmylated T3, a 54nt RNA, about half as long as the 100nt RNA. One obvious difference between the two profiles is the relative absence of bumping events from the T1 *i-t* trace, perhaps indicating an additional way that the pore discriminates oligos based on length; more experiments needed to clarify this point.

To probe (b) two 32nt oligos were designed with 9 or 13 pyrimidines, lined in groups with up to 5 adjacent pyrimidines (see T7(13) and T8(9) in Table 1); please note that the 3’end is identical to the sequence most abundant in tRNA and currently in some sgRNA. Evidence that adjacent pyrimidines are fully osmylated using Yenos’ protocol, is shown by the agreement between observed and theoretical R(312/272) values; see Table 1 and discussion above. Experiments with these oligos were conducted with a biased voltage at −220mV, and/or *I*_*o*_ ≈ 270pA, considered sufficient to induce numerous translocations (see Figures S13 and S14 in the Supplementary Information, and discussion below). Noteworthy is that the MinION can successfully run experiments with applied voltage as high as at −220mV.

A fourth question that was not fully investigated is whether the MinION may be used for quantitation of osmylated RNAs. The experiments reported here are conducted with a formal oligo concentration in the 1 to 2 µM range in the ONT buffer, typically obtained by dilution in the buffer from an RNA(OsBp) stock solution of 20 to 40 µM. Considering that the pyrimidine osmylation is practically 100% and that purification leads to practically full recovery with no dilution, it will be of interest to quantify RNA(OsBp) by nanopore and deduce the concentration of a certain RNA in an unknown sample. Limited experiments with dilutions in the range of 3-to 10-fold from the typical concentration illustrate proportionality between concentration and number of actual translocations and supports quantitation. To further probe this issue the software, buffer, and pore temperature, which is currently at 34°C, of this platform need to be optimized for unassisted translocation of osmylated RNA.

Over a period of 10 months a total of 8 different flow cells was used, primarily due to our inexperience with this platform and the novelty of the application. Over 70 experiments were conducted lasting less than an hour each. All the experiments reported here were conducted with flow cells containing 512 channels. Recently ONT commercialized the Flongel that contains 50 channels; Flongel should be a more suitable flow cell for many applications including the one described here.

### Acquisition of MinION data, reporting, and results

The software of the MinION, MinKNOW, is currently set up for sequencing experiments, and not for analysis of single molecule translocation data. A recent MinKNOW update includes acquisition of raw *i-t* recordings in *fast-5* file format and direct visualization with MatLab software on a *i-t* plot that includes a units grid, which allows one to precisely determine *I*_*o*_, *I*_*r*_ and *τ* for any event at an apparent accuracy much better than any instrument could deliver. This feature permits two people, one reading and one reporting, to estimate and report lowest *I*_*r*_ values for every event at ±(1 to 2) pA at a rate of about 700 data per hour. Concatenated *i-t* traces such as the ones reported in Figure 4 were obtained in house by saving the *fast-5* files in *txt* format readable by open source software QuB (see Experimental Section) and manually “cleaning up” the file to remove instrument lines as well as a lot of events that can’t be attributed to single molecule translocations, as it will be described next.

Close inspection of the recordings revealed that events, with the exception of the above mentioned “instrument lines” (Figures 5A and 5B), could be grouped into four categories: (i) Highly noisy events with randomly variable *I*_*r*_, and duration in the range of a few seconds that seem to increase with voltage, oligo concentration, and flow cell use; these events were attributed to noise and ignored. (ii) Relatively shallow and short, 1 to 2 ms long, events that measure 0.4 < *I*_*r*_*/I*_*o*_ < 0.9, presumably resulting from molecules bumping at the pore were also ignored. (iii) Long events with *I*_*r*_*/I*_*o*_ < 0.4 that appear to be the result of multiple translocations, one molecule following the other without reaching open pore current between events; these were also ignored. (iv) Last, but not least, events were observed of low noise level with *I*_*r*_*/I*_*o*_ < 0.4 and durations in the range of 2 to 200 milliseconds. Such events account for about 150 to 250 (at about 2µM oligo concentration) per half an hour of *i-t* recording depending on molecule length. It is estimated that this count should be at least 5-times higher and approach 1000 translocations per half an hour, if one were to count in molecules that translocate as part of group iii, or if the system is optimized so that group iii is suppressed. Please note that with molecules in group iii the observed *I*_*r*_ is likely not comparable to the corresponding *I*_*r*_ for single molecule translocation due to occupation of the pore by more than one molecule at the time. We attributed group (iv) to single molecule translocations and reported the lowest observed *I*_*r*_ (pA) value for each translocation. Initially we also reported duration (*τ*) for each translocation (Figure S19, Supplementary Information), but this type of information, due to the stochastic nature of the process, did not appear to provide added insight for our current application. Experiments with 31nt osmylated T11, T12, T14 and T15 (see Table 1), conducted at both −140mV and −180mV, strongly suggest that mean durations of translocations decrease with increasing voltage, confirming that the events of group iv are true translocations.

Values *I*_*o*_ and *I*_*r*_ were estimated from the *i-t* traces in the Matlab plots, reported together from different experiments and from different channels of the same experiment in Microsoft Excel, *I*_*r*_*/I*_*o*_ calculated from the corresponding *I*_*o*_, and *I*_*r*_*/I*_*o*_ graphed as a histogram with bin=0.01, 0.02 or 0.04 depending on the oligo; see histograms for each tested RNA in the corresponding Supplementary Information figure. Histograms of two structurally similar RNAs are compared (or fingerprinted) against each other by normalizing the counts of each bin for the total number of translocations as seen in Figure 6.

### I_r_/I_o_ histogram is the fingerprint of an RNA(OsBp)

Each figure in Figure 6 compares RNAs (Table 1) that differ minimally from each other and illustrates cases where the nanopore senses, or not, small structural changes on the tagged pyrimidine moiety, or on the adjacent nucleobase, presumed to be the base following and not the base ahead of the osmylated moiety. Figure 6A compares translocation fingerprints of osmylated T11(AC*A) and T12(AU*A); for simplicity OsBp is indicated with a star (*). As a reminder the difference between C and U is that C contains an NH_2_ moiety on C4, whereas U contains an Oxygen (O) on C4, hence the mass unit difference between these two RNAs is 1/10,500, where 1 mass unit is the difference between NH_2_ and O and about 10,500 is the molecular weight of a singly osmylated 31nt RNA. Figure 6A illustrates that the *I*_*r*_*/I*_*o*_ profile (fingerprint) of T11(AC*A) has, perhaps, up to four maxima with (*I*_*r*_*/I*_*o*_)_max_=0.08, 0.15, 0.19 and 0.23, whereas the *I*_*r*_*/I*_*o*_ profile of T12(AU*A) has two maxima at (*I*_*r*_*/I*_*o*_)_max_=0.12 and 0.17. One may presume that the translocation(s) with the least residual ion current corresponds to a 3’-entry and the one with the higher residual ion current corresponds to 5’-entry, in analogy to the observations with intact nucleic acids.^53^ T11(AC*A) appears to incorporate a total of four (*I*_*r*_*/I*_*o*_)_max_ to correspond, tentatively, to the four different configurations of interaction between a singly osmylated RNA and the nanopore (see Figure 3C and discussion above). These four possible configurations between RNA(OsBp) and nanopore may or may not yield distinct or detectable *I*_*r*_*/I*_*o*_ levels, as indicated by the fewer than four (*I*_*r*_*/I*_*o*_)_max_ values observed with most of the 31nt oligos. For the purpose of the following discussion the conjecture is made that, if a motor enzyme could be engineered to process osmylated RNA one base at a time, then the observed (*I*_*r*_*/I*_*o*_)_max_ values from the unassisted translocations will correspond closely to the values to be observed under enzyme-assisted translocation. Figure 6A illustrates a 4% difference (from 0.08 to 0.12 pA, i.e. between the two lowest (*I*_*r*_*/I*_*o*_)_max_ values) that, for a typical *I*_*o*_ =200 pA, translates to 8pA. A difference of 8pA suggests that A-C*-A subsequence is well discriminated from the A-U*-A in this platform, indicating that the above envisioned enzyme-assisted translocation, would result to a highly accurate base calling when C* or U* is followed by A, located within an oligoadenylate sequence, to be precise. Assuming that the nucleotide ahead of the tagged one (A here), may not contribute much to the observed discrimination, we propose that this platform senses a two nucleotide subsequence, when the first nucleotide is tagged.

Figure 6B compares normalized histograms *I*_*r*_*/I*_*o*_ from the experiments with T11(AC*A) and T14(GC*G), and illustrates the effect of replacing the adenosine with guanosine. It appears that (*I*_*r*_*/I*_*o*_)_max_=0.08 and 0.19 with AC*A are now shifted to more residual ion current (*I*_*r*_*/I*_*o*_)_max_=0.13 and 0.21 with GC*G. This is a major shift and supports the above proposition of sensing a two nucleotide subsequence, when the tagged nucleotide is Cytosine. In contrast to the effect observed with tagged C, tagged U doesn’t result to such large effect, i.e., yielding comparable *I*_*r*_*/I*_*o*_ profiles between T12(AU*A) and T15(GU*G) (Figure 6C); hence the corresponding (*I*_*r*_*/I*_*o*_)_max_=0.12 and (*I*_*r*_*/I*_*o*_)_max_=0.13 may yield a 2pA (from (0.13-0.12)*200pA) distinction only. Enhanced discrimination in this case may be achieved in the presence of a second tag to react selectively with Guanosine and leave the other bases intact.

In contrast to Figure 6A, Figure 6D illustrates practically identical fingerprints for osmylated T13(A-5-MeC*-A) and T16(A-5-MeU*-A), pointing out that the presence of a methyl group reduces the ion current so much ((*I*_*r*_*/I*_*o*_)_max_ = 0.03), that the difference in the pyrimidine moiety becomes “silent”. Whether or not replacing A with G adjacent to the 5-Me-pyrimidine will make a difference, remains to be tested. It is noticeable that replacing a canonical pyrimidine with its 5-Me derivative yields 10pA for C (from (0.08-0.03)*200pA) and 18pA for U (from (0.012-0.03)*200pA) lower maximal residual ion current leading to, perhaps, the most accurate base calling envisioned for 5-Me modifications.

Since the specific nanopore platform does not discriminate 5-MeC from 5-MeU, one could use the higher reactivity of OsBp for 5-MeU over 5-MeC, C, U, and 4-SU (compare part B in Figures S1-S7 in the Supplementary Information), and prepare RNA(OsBp) at low OsBp concentration, where mostly 5-MeU moieties are osmylated and the other pyrimidines are not. This would be an example where the selectivity of the pyrimidine-specific label assists in discrimination, when the nanopore falls short.

All but one the tested 31nt RNAs exhibited their lowest (*I*_*r*_*/I*_*o*_)_max_ in the range of 0.03 ≤ (*I*_*r*_*/I*_*o*_)_max_ ≤ 0.13. This range falls below the (*I*_*r*_*/I*_*o*_)_max_ = 0.15 reported for the intact 74nt T6, and far below the (*I*_*r*_*/I*_*o*_)_max_ = 0.31 observed with poly(I) (Figure 6E); see discussion later. The outlier is the 31nt T17(A-4-SU*-A) which exhibits (*I*_*r*_*/I*_*o*_)_max_ = 0.27 (Figure 6E). One explanation for the observed high (*I*_*r*_*/I*_*o*_)_max_ = 0.27 could have been that osmylation of T17 may have led to phosphodiester cleavage at the 4-SU base, and that osmylated 15nt or 16nt is the main translocating material. This explanation is excluded based on HPLC analysis of the osmylated product with an IEX method known to yield resolution based on oligo length^49^ (see Experimental Section); the analysis reveals no detectable shorter oligos. The osmylation mechanism in the presence of tertiary nitrogen donor ligands, like bipy, as presented in Figure 2A is well documented.^56^ An explanation based on the reactivity of OsO_4_ towards uracil to form cytosine^57^ is not applicable under our conditions, since there is no ammonia present in the OsBp reagent and therefore transformation of C=S to C-NH_2_ is impossible. Moreover T11(A-C*-A), which would have been the product of this transformation, translocates with (*I*_*r*_*/I*_*o*_)_max_ = 0.08 and not 0.27 (Table 1). Another explanation based on the reactivity of a catalytic amount of OsO_4_ in the presence of reoxidants, to convert alkenes into cis-vicinal diols is also not applicable^58^ as OsBp contains no reoxidants and diol formation with concurrent OsO_4_ removal will not exhibit absorbance at 312nm, in contrast to the observations (Figure S7 parts B. & C.). CE analysis of the osmylated T17, as well as the osmylation kinetics monitored by CE using the long capillary indicated the formation of two separate products with two topoisomers each (Fig S7 in the Supplementary Information), all four products are stable at room temperature under extended osmylation conditions, suggesting reaction of OsBp at two locations on the 4-SU moiety, perhaps a C4-C5 conjugation in addition to the typical C5-C6 conjugation, consistent with Diode Array spectroscopic data. Clarity on this issue may await further experimentation, and the current rationalization for the nanopore translocation features of this outlier is attributed to desolvation, as will be described later.

### Identification of RNA oligos by HPLC vs. nanopore

Discrimination among the seven 31nt RNAs, due to their structural similarity in being practically oligoadenylates, is challenging. Osmylation broadens the oligo peak, or splits it in two due to the presence of two topoisomers, and therefore HPLC analysis of osmylated oligos will not yield better resolution. A comparison can be made between resolution of the intact oligos by HPLC and discrimination of the osmylated oligos by nanopore. In this context only T17 is easily discriminated from the others by both HPLC and the MinION (compare Figure 3A with (*I*_*r*_*/I*_*o*_)_max_ data in Table 1). Histograms of osmylated T13 and T16 (Figure 6D) are identical, histograms of osmylated T12 and T15 are quite similar (Figure 6C), but histograms of the other three oligos are distinct. In contrast the best HPLC method available to us discriminates between two oligos but yields no baseline resolution among the other four. In addition, resolution by HPLC is reduced as a function of the oligo length, meaning that if the A15nt tails were to be replaced by A25nt tails, then HPLC resolution will become undetectable. In contrast, a nanopore interrogates the molecule as it passes through, and length is practically a non-issue, meaning that extending the tails from A15nt to A25nt is not expected to diminish discrimination. Hence it appears that nanopore characterization of RNA(OsBp) is in certain cases superior to intact RNA analysis by HPLC.

Figure 6F compares normalized *I*_*r*_*/I*_*o*_ histograms from two 22nt oligos, i.e. T31(AC*C*A) and T32(AU*U*A). Comparing Figures 6A,B,C with 6F illustrates that two adjacent Py(OsBp) produce measurably less residual ion current compared to one Py(OsBp) followed by a purine, substantiating our argument for two-base discrimination in this system. In agreement with Figure 6A that illustrates less residual ion current for C compared to U, Figure 6F illustrates that two adjacent osmylated Cs yield less residual ion current compared to two adjacent osmylated Us, with (*I*_*r*_*/I*_*o*_)_max_ = 0.03 and 0.06 for T31 and T32, respectively. If any, the shorter sequences should have produced more, not less, residual ion current, but the length appears to play a smaller role here. The shorter sequences (22nt vs 31nt) were selected for better resolution of the three diastereomers obtained by osmylation as monitored by CE (see Figure 3B). The smaller role of the length with unassisted RNA(OsBp) translocation is consistent with the histograms comparison in Figure 6G between osmylated 53nt T3(UU-UU) and 74nt T6(UU-UU-UU), that contain U*U* separated by a A_19_ subsequence. Specifically (*I*_*r*_*/I*_*o*_)_max_ = 0.03 for both, while the profile of the longer oligo exhibits relatively more counts towards less residual ion current compared to the shorter oligo. This feature can be rationalized on statistical grounds considering that molecules with two U*U* in OL configurations (see Figure 3C) are more abundant with the long vs the short oligo. It is noticeable that (*I*_*r*_*/I*_*o*_)_max_ with T3 and T6 measures 0.03, whereas (*I*_*r*_*/I*_*o*_)_max_ with T32(UU) measures 0.06, indicating that the residual ion current inside the pore for unassisted translocations is modulated by the presence of a sequence of, at least, 4+19=23 nucleotides (from UU+UU+19As). This conclusion is strictly valid for an unassisted translocation that yields a single event, and not applicable to either motor-enzyme assisted translocations or to translocations that yield ion current modulation dependent on base sequence, as shown by inter-event detail (see later discussion). Figure 6H compares normalized histograms of osmylated T7(13Py) and T8(9Py), indicates (*I*_*r*_*/I*_*o*_)_max_ =0.01 for both, and illustrates closely similar profiles with minimal differences suggesting that during unassisted translocation a certain number of OsBp moieties, perhaps 9 or less per total of 32nt, is responsible for obstruction and the rest of the moieties are not making a difference, implying that obstruction of ion current due to the OsBp presence may not be as dramatic as one were to assume based on size. These two oligos in addition to the 100nt sgRNA were specifically designed with five adjacent pyrimidines at the 3’-end and were tested in order to cofirm that t-RNAs and sgRNAs will translocate via the MinION. Last but not least Figure 6I compares the normalized histograms of the two 22nt miRNAs^59^ and illustrates some differences that appear to be outside experimental deviation. miRNA122 has a three pyrimidine sequence at the 3’end and single pyrimidines spread through the sequence, whereas miRNA140 has mostly purines at the ends of the strand and most of the pyrimidines lined in the middle of the sequence. These two miRNAs could, in principle, be discriminated based on their different histogram profiles, but it could be more useful, if there is additional information to discriminate them within a mixture of miRNAs.

### Is there characterization/sequencing information in unassisted RNA(OsBp) translocations?

The fingerprints shown in Figure 6 are not the only information obtained from this platform. They are definitely helpful in identifying one from another in certain cases. However the real value would be if one could identify and quantify each in a mixture. For example, (i) a nanopore-based assay to identify a panel of miRNAs from a blood or urine sample^59-60^ will find application in medical diagnostics or (ii) a nanopore-based assay to determine purity and impurities in a sample of sgRNA will find application in pharmaceutical development.^49^ The first question to ask is whether nanopore-based analysis has such potential. Let us consider a platform improved over the one explored here, so that there are not only 250, but about 1000 single molecule translocations per half-an-hour of experimentation (see above). Considering the OsBp isomerism in producing several diastereomers, and selecting only for the ones that have the lowest possible *I*_*r*_ which is attributed to PA or OL configurations (see Figure 3C), one may end up with a mere 25% or 250 translocations out of the 1000. Assuming that only 20% of those are of long enough duration, let us say longer than 10ms and exhibit inter-event detail, then only 50 of those may prove “useful”. Hence each channel may provide 50 translocation with sequence-bearing information per 30min data acquisition. If only 75% out of the 512 channels in a MinION flow cell are in good working condition, that leaves 384 working channels and yields 50*384=19,200 sequence-information bearing events. Assuming a sample with 100 miRNAs, there is on average a 192-fold representation of each miRNA in the sample, more or less depending on their concentration. Evidently an experiment can last multiples of 30min, or let us say 2.5 hours, and produce a total of about 100,000 useful events to identify and quantify 100 miRNAs. This “Gedanken experiment” is rather promising, considering that it forgoes improvements in sampling rate and/or label. The case in favor of implementing a nanopore-based 100nt sgRNA purity assay is even more appealing, as one expects that the impurities may be less than 100.

Since the potential exists, we present below a practical approach of how to obtain sequence-bearing translocations and show such examples in Figures 7 and 8. Initially one, or even better an algorithm, has to identify translocations of group iv (see above), then identify the ones from group iv that have dwell times over 10ms, and then select the ones that produce the lowest *I*_*r*_. Expected value(s) for the lowest *I*_*r*_ for a certain molecule can be approximated using the (*I*_*r*_*/I*_*o*_)_max_ listed in Table 1. This process ignores single molecule translocations with AP configurations of adjacent pyrimidines and simplifies the selection. Following this approach we visually identified the examples shown in Figures 7 and 8. Figure 7A compares translocations obtained at −180mV from osmylated T6 (left) and intact T6 (right). Observed dips at lower *I*_*r*_ levels (OL for the lowest and PA for the other two) are attributed to the three U*U* and the red dotted line illustrates the *I*_*r*_ level of the adenosines in between. Figure 7B presents examples of T4(CC-UU) translocations at −180mV obtained from different channels, exhibiting different dwell times τ, but with the same number (2 “dips”, pink blocks), of low *I*_*r*_ levels, attributed to Py*Py*, separated by a higher *I*_*r*_ level attributed to Pu (dotted red line). Sequence of T4 is listed in Table 1 and the histogram is in Figure S11 in the Supplementary Information; notably a 10-fold lower concentration (0.1 µM) used for this experiment led to about a 10-fold decrease in translocations. Figure 7C presents the best molecule in this study to highlight rudimentary sequencing as it contains three heavily osmylated pyrimidine sequences separated by purine sequences. Examples of T7(13Py) translocations obtained at −220mV from three different channels illustrates the expected three dips of low *I*_*r*_ level that correspond to the three osmylated sequences of consecutive pyrimidines and the higher *I*_*r*_ level that corresponds to the purines in between; notably purine sequence in T7 is not the same as in T6.

**Figure 7.**
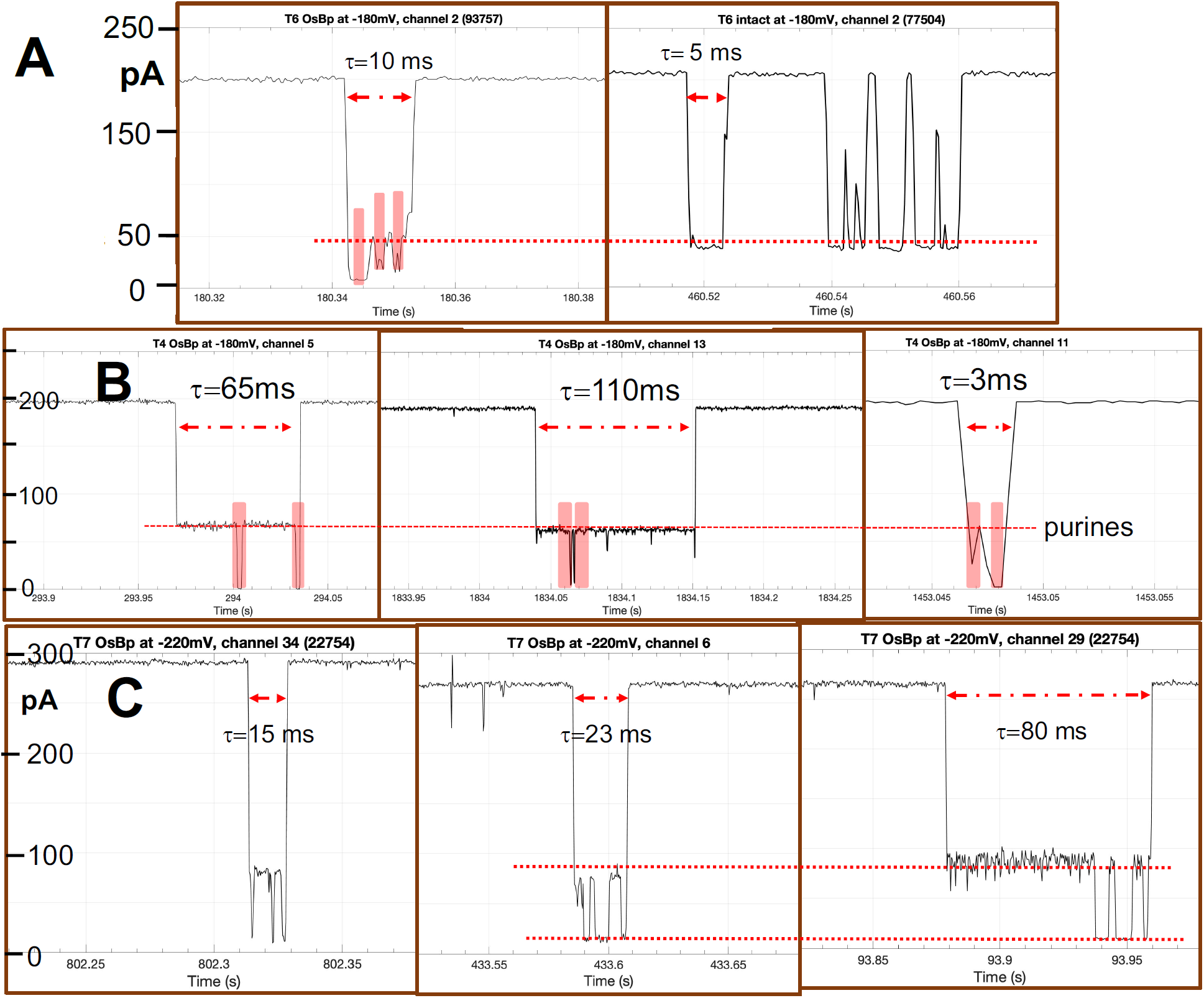
**Inter-event detail to suggest rudimentary “sequencing”,** as shown by the different *I*_*r*_ levels within a single molecule translocation, attributed to Py(OsBp)_n_ or Pu_m_, where n and m are the number of adjacent Py and Pu, respectively. All presented molecules are practically 100% pyrimidine-osmylated. **A.** Comparison of T6 translocations at −180mV between the osmylated (left) and the intact (right) oligo. Observed dips at low *I*_*r*_ levels to correspond to the three osmylated U doublets and the red dotted line illustrates the *I*_*r*_ level of the adenosines in between. **B.** Examples of T4(CC-UU) translocations at −180mV obtained from different channels, exhibiting different dwell times *τ*, but with the same number, i.e. two “dips” (pink blocks), of low *I*_*r*_ levels, attributed to Py(OsBp)Py(OsBp), separated by a higher *I*_*r*_ level attributed to Pu (dotted red line). Noteworthy that the *I*_*r*_ level attributed to the consecutive pair of pyrimidines could be one of three, consistent with the three different diastereomers (see Figures 3B and 3C). **C.** Examples of T7(13Py) obtained at −220mV from three different channels illustrates the expected three dips of low *I*_*r*_ level to correspond to the three osmylated sequences of consecutive pyrimidines and the higher *I*_*r*_ level that corresponds to the purines in between.

**Figure 8.**
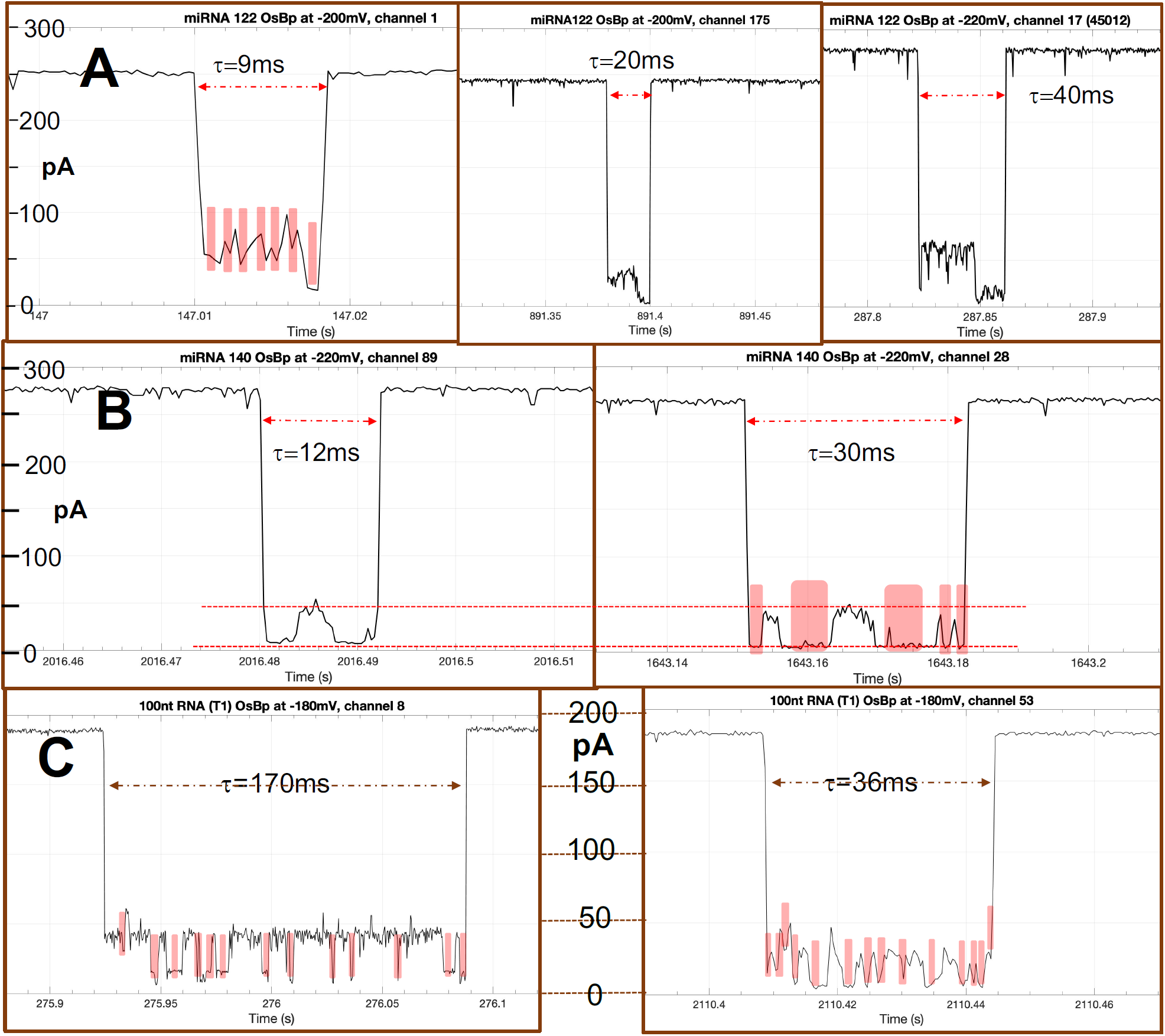
**Inter-event detail to suggest rudimentary “sequencing” of miRNAs and sgRNA**, as shown by the different *I*_*r*_ levels within a single molecule translocation, attributed to Py(OsBp)_n_ or Pu_m_, where n and m are the number of adjacent Py and Pu, respectively. All presented molecules are practically 100% pyrimidine-osmylated. **A.** Three examples of miRNA122 translocations taken from two different experiments (left and middle at −200mV, right at −220mV); all show translocations consistent with entry from the 5’-end consistent with higher *I*_*r*_ level that corresponds to single pyrimidines interspersed among purines, and exit from the 3’-end consistent with low *I*_*r*_ level that corresponds to a sequence of 3 consecutive pyrimidines. The left figure highlights with pink blocks the dips that may be attributed to the six single Py(OsBp), all six at a higher *I*_*r*_ level compared to the *I*_*r*_ level of the last dip that should correspond to the three consecutive Py(OsBp), i.e. 5’ **U**GG AG**U** G**U**G A**C**A A**U**G G**U**G **UUU** G 3’. **B.** Two examples of miRNA140 translocations taken from two different channels of experiment conducted at −220mV. Red dotted lines illustrate the higher *I*_*r*_ level attributed to the purines, and the low *I*_*r*_ level attributed to the two subsequences with consecutive Py(OsBp). The translocation with the longer duration (right) appears to match closely the sequence of miRNA140, in the form of Py followed by Pu, i.e. 5’ **C**AG **U**GG **UUU U**A**C CCU** A**U**G G**U**A G 3’. **C.** Two examples of the 100nt sgRNA, T1, obtained at −180mV from two different channels with 14 dips (pink blocks), consistent with the 14 subsequences of this RNA that contain two or more Py (highlighted in red below). Noteworthy is the better resolution between purine and pyrimidine regions observed with the translocation that exhibits the longest duration of 170ms. The five consecutive pyrimidines at the 3’end should have produced a much deeper dip, but at −180mV it is likely that only a truncated version of T1, let us say one missing the last 2 or 3 nucleotides, translocated. T1 sequence, 14 red highlighted regions with 2 or more consecutive Py: 5’-**UU**A CAG **CC**A CG**U CU**A CAG CAG **UUU U**AG AG**C U**AG AAA UAG CAA G**UU** AAA AUA AGG **CU**A G**UC C**G**U U**A**U C**AA **CUU** GAA AAA GUG GCA **CC**G AG**U C**GG UG**C UUU U**-3.

Last but not least, Figure 8 illustrates the potential of this approach for unassisted characterization of a mixture of miRNAs (Figures 8A and B) and for a synthetic preparation of sgRNAs (Figure 8C). Figure 8A presents three examples of miRNA122 translocations taken from two different experiments (left and middle at −200mV, right at −220mV); all show translocations consistent with entry from the 5’-end, as shown by the higher *I*_*r*_ level that corresponds to single Py* interspersed among purines, and exit from the 3’-end consistent with low *I*_*r*_ level that corresponds to a sequence of 3 consecutive pyrimidines. The left figure highlights with pink blocks the dips that may be attributed to the six single Py* separated by purines (see sequence in Table 1), all six at a higher *I*_*r*_ level compared to the *I*_*r*_ level of the last dip that is attributed to the three consecutive Py* at the 3’end. Figure 8B presents two examples of miRNA140 translocations taken from two different channels of an experiment conducted at −220mV. Red dotted lines illustrate the higher *I*_*r*_ level attributed to the purines, and the low *I*_*r*_ level attributed to the two subsequences each with four consecutive Py*. The translocation with the longer duration (right) appears to match closely the sequence of miRNA140, in the form of Py* followed by Pu. Figure 8C presents two examples of 100nt sgRNA (T1, see sequence in Table 1) translocations obtained at −180mV from two different channels with 14 dips (pink blocks), consistent with the 14 subsequences of this RNA that contain two or more Py(OsBp). It is noteworthy that the better resolution between purine and pyrimidine regions is observed with the translocation that exhibits the longer duration of 170ms. The examples in Figures 7 and 8 suggest that the longer translocations, e.g. in the range of 10 to 200 ms carry inter-event detail reflecting the sequence of Py(OsBp) separated by a sequence of intact purines. Hence optimization of the platform conditions towards single molecule characterization is worth pursuing. It is also conceivable that a processing enzyme could be produced by directed-molecular evolution to work with osmylated RNAs or any other type of tagged RNAs, process them one-base at a time, and “sequence” them via a two-base discrimination such as the one observed in this study.

### Parameters that affect residual ion current in the MinION platform

The data reported here are obtained with the MinION, and may be only partially applicable to other nanopore platforms. When we initiated the project of combining nucleic acid labeling with nanopore single molecule analysis, we were under the impression that the bulkiness of OsBp will dictate nanopore size-suitability. The first surprise came by observing translocations of osmylated oligos via α-HL, which has been the prototype protein pore for intact nucleic acids. How is it possible that the increased bulkiness of RNA(OsBp) fits via the same pore size as intact RNA? The second surprise was observing translocations of T7 and T8 (both with a five consecutive Py(OsBp) subsequence) via the MinION pore protein, albeit at the higher biased voltage of −220mV. How is it possible to use a platform suitable for intact RNA and detect translocations of overlapping OsBp moieties? The third observation, reported 30 years ago with *α*-HL, and confirmed experimentally here with the MinION platform, are the relatively low *I*_*r*_ levels observed with the intact homopolymers poly(C) and poly(U). The fourth observation is the relatively high *I*_*r*_ observed for oligoadenylates (*I*_*r*_*/I*_*o*_ =0.15 here), opposite to expectation considering that sterically adenine is about double the size of a pyrimidine. In addition to the already proposed rationalizations for the last two observations that attribute these phenomena to homopolymer special helical structures in a confined environment,^1,2^ we wish to propose hydration or solvation/desolvation, as another critical parameter, and envision this phenomenon as follows: There are two different sources of water molecules present within the nanopore at all times. One source are the water molecules solvating the salt ions which are responsible for the observed ion current, and the other source are the water molecules solvating exposed functional groups of the amino acids of the protein pore as well as solvating functional groups of the translocating nucleic acid. Within the confined space of the pore less water molecules used for solvation of the pore protein and the translocating nucleic acid directly translates to more water molecules available to transfer salt ions through the pore, or the equivalent, i.e. more ion current. Solvation inside a nanopore will not resemble solvation in the bulk solution, but desolvation is costly from a thermodynamic point of view and the first, perhaps the second too, solvation shells of any functional group within the pore will remain intact, unless it is replaced by a new component in the system. In this context higher solvation requirements can rationalize the typically lower *I*_*r*_ of ribooligos compared to deoxyoligos, the typically lower *I*_*r*_ of pyrimidines (two functional groups) compared to adenine (one functional group), and the reported here high *I*_*r*_ level of 4-SU moiety compared to a U base, based on the known lower solvation needs of C=S *vs* C=O. Reduced solvation requirements may also rationalize the apparently lessened effect of OsBp’s size by considering that OsBp moiety with its almost parallel to the strand backbone configuration serves as a “shield” and by steric hindrance replaces water molecules that otherwise accompany the nucleic acid. “Touching” proximity between tagged-RNA and nanopore leads to desolvation and reorganization that yields superior recognition and discrimination. A lot remains to be learned from the interactions between labeled RNAs and nanopores, and exploitation of this novel approach appears promising for analytical assays in pharmaceutical applications and for small RNA diagnostic assays.

## CONCLUSIONS

Earlier nanopore studies with intact DNA/RNA exploited an immobilized strand within a nanopore in order to determine sequence-dependent *I*_*r*_*/I*_*o*_ values. Translocation of osmylated RNA oligos is dramatically slower, even in the presence of one OsBp moiety per oligo, the residual ion current is markedly reduced, and unassisted, voltage-driven translocation illustrates recognition and discrimination of minimal chemical structural changes, such as an C-NH_2_ group in place of a C=O, or an additional methyl group on the pyrimidine moiety, or one pyrimidine followed by another pyrimidine instead of followed by a purine. The platform that showcased these fine recognition traits is commercially available, easy to use, relatively inexpensive, and could be further optimized for the characterization and identification of short RNAs in mixtures. This study illustrates the contribution of the labeling approach for improved recognition. RNAs in the range of 22 to 300nt have important functions and are currently promoted as pharmaceuticals. A simple assay to assess identity and quantity of miRNAs, as a non-invasive diagnostic test, could revolutionize patient care and assess health status for all individuals. Critical human microRNAs are believed to be about hundred, can be found in most biological fluids, and an assay that yields Py/Pu sequence to identify most of them, may be easy to develop, implement, and commercialize.

## EXPERIMENTAL SECTION

### Materials and Methods

#### RNAs and other Reagents

Custom-made RNA oligos were purchased from Trilink Biotechnologies or Dharmacon (Horizon Discovery Group); their sequences and the properties of their osmylated derivatives listed in Table 1. Custom-made deoxyoligos were purchased from Integrated DNA Technologies (IDT). mRNA Cas9 and mRNA EGFP (both CleanCap) were purchased from Trilink Biotechnologies. tRNA(Cys), the E.Coli version by in-vitro transcription was purchased from tRNA probes, Texas. Homopolymers poly(C), poly(U) and poly(I) were purchased from Sigma-Aldrich. Purity of oligos was tested by HPLC in-house, typically at or below 85%, but varies depending on the purification level that was requested from the manufacturer. RNAs were diluted with Ambion Nuclease-free water, not DEPC treated, from Thermo Fisher Scientific typically to 200 or 400 µM stock solutions and stored at −20°C.

Buffers DNase- and RNAse-free TRIS.HCl 1.0 M pH 8.0 Ultrapure was purchased from Invitrogen. and 1.0M pH 7 from Sigma. Triethylamine acetate buffer 2.0M pH 7 and Sodium hydroxide 1.0M bioreagent were purchased from Sigma-Aldrich; KCl and NaCl crystalline ACS min 99.0% from Alfa Aesar. 50mM sodium tetraborate buffer pH 9.3, HPCE quality, from Agilent Technologies. Wt alpha Hemolysin (a-HL) monomer was purchased from List Biologicals, Mountain View, CA. Distilled water from ArrowHead or Alhambra was used for preparation of HPLC mobile phase. A 4% aqueous osmium tetroxide solution (0.1575 M OsO_4_ in ampules at 2 mL each) was purchased from Electron Microscopy Sciences. 2,2’-Bipyridine 99+% (bipy) was purchased from Acros Organics.

#### Osmylation and purification

OsBp reagent was prepared by preweighing the equivalent of 31.4 mM of 2,2’-bipyridine (bipy) in 8 mL of water in a scintillation vial and adding the full content (2mL of a 4% OsO4) supplied in an ampule in order to prepare a 10mL 31.4 mM OsBp stock solution, 1:1 in OsO_4_ and bipy. The concentration of the OsBp stock solution is limited by the solubility of bipy in water and adding OsO_4_ does not increase it, as the complex has a low association constant; OsBp complex represents an approximate 5% of the total, as measured by CE^41^. The low association constant of this complex is also consistent with the dependence of the observed osmylation rate on the square of the nominal concentration [OsBp] (see Table S1 and Figure S17 in the Supplementary Information). Care should be taken that this preparation is conducted in a well ventilated area and that all leftover traces of OsO_4_ are properly discarded. The freshly prepared stock solution is then dispensed in HPLC vials and kept at −20°C; each vial can be stored at 4°C and used for a few weeks without loss of potency. It is recommended that every separate stock solution is being validated before first use. To ensure that pseudo-first order kinetics apply we typically use an excess of OsBp at 25-fold or larger compared to the reactive pyrimidine in monomer equivalents. Typically the reactivity of the mononucleotide mirrors the reactivity of this base within an oligo. Manufacturing of osmylated RNAs was conducted at about 12mM OsBp, and purification from excess OsBp was done with spin columns (TC-100 FC from TrimGen Corporation) according to the manufacturer’s instructions which takes about 8 min. Close to 100% recovery of RNA is achieved with minor volume/concentration changes, and OsBp reagent is reduced to undetectable amounts after 2xpurification.

#### HPLC and CE methods

Analyses targeting purity and resolution of oligos of similar sequence were conducted by gradient HPLC; both IEX and IP-RP modes were exploited. Kinetic measurements were primarily conducted by CE, that requires less analysis time compared to gradient HPLC, and therefore relatively faster reactions can be monitored better by CE. All analyses were conducted automatically; autosamplers are thermostatted and samples were in queue kept at the reported temperature. Both CE and HPLC peaks were detected and identified using a diode array detector (DAD) in the UV–vis region 200–450 nm and the electropherograms or chromatograms were recorded at 260, 272 and 312 nm; they are reported here selectively. Samples were prepared with RNAse free water, but buffers were not. No RNA degradation has been observed in our Laboratory.

For HPLC analysis we used an Agilent 1100/1200 LC HPLC equipped with a binary pump, Diode Array Detector (DAD), a 1290 Infinity Autosampler/Thermostat, and Chemstation software Rev.B.04.01 SP1 for data acquisition and processing. As IEX HPLC column DNAPac PA200 from ThermoFisher Scientific (Dionex) was used in 2X250 mm or 4×250mm configurations. The performance of the instrument and the column was qualified using standards every time ahead as well as after analysis of research samples. Two IEX HPLC methods were routinely used and have been validated for purity determination and RNA stability. IEX method at pH 8 is exploiting a 1.5M NaCl gradient in a 25mM TRIS.HCl pH 8 buffer, with 30°C column compartment; typical gradient 15%B to 55%B in 12min where A is 25mM TRIS.HCl and B is 1.5M NaCl in A. IEX method at pH 12 is exploiting a 1.5M NaCl gradient in a 0.01 N NaOH solution (no other buffer needed) with 10°C column compartment; typical gradient is 0%B to 95%B in 16 min where A is 0.01M NaOH and B is 1.5M NaCl in A. IEX method pH 12 is validated by us and is recommended for longer RNAs in order to suppress secondary structure that broadens peaks and yields low and misleading resolution^49^. An IP-RP HPLC method was also employed to test RNA resolution with HPLC column DNAPac RP from ThermoFisher Scientific (Dionex) in 2X100 mm configuration and flow at 0.35mL/min. Method IP-RP is exploiting a 25%v/v acetonitrile-water gradient in a 0.1 M TEAA buffer pH 7 at 30°C column compartment^49^.

CE measurements were conducted with an Agilent G1600 Capillary Electrophoresis (CE) instrument equipped with DAD and Chemstation software Rev.B.04.03(16) for data acquisition and processing; the CE was used in conjunction with a circulating bath to control the autosampler’s temperature. The capillary’s temperature was controlled by the instrument’s software. Typical capillary zone electrophoresis (CZE) analyses were conducted with an untreated fused silica capillary (50 µm × 40 cm), coined “short” capillary in pH 9.3 50 mM sodium tetraborate buffer using 20 kV or 25 kV. Same buffer was used with an untreated fused silica capillary (50 µm × 104 cm), coined “long” capillary to resolve the topoisomers from the osmylated oligos (Figure 3B and Figures S1-S7, S13, S14, Part C). Both capillaries, short and long, were equipped with an extended light path and were purchased from Agilent Technologies.

#### Single molecule translocations with wt α-HL/EBS GNM membrane (EBS platform)

A limited number of experiments was conducted with the wt a-HL/EBS membrane (EBS) platform (for examples, see Figure S18 in the Supplementary Information). Nanopore experiments were conducted with 10 µM synthetic DNA/RNA oligo in 1.0 M KCl, 10 mM TRIS.HCl buffer at pH 8.0 and at 20 ± 1 °C, as described in detail^61^, and summarized here. wt α-HL was purchased from List Biological Laboratories in the monomer form of lyophilized power and dissolved in water at 1 mg/mL. 1,2-Diphytanoyl-sn-glycero-3-phosphocholine (DPhPC), kindly provided to us from EBS, was dissolved in decane at 10 mg/mL and used to form the bilayer. The bilayer was supported by a glass nanopore membrane (GNM) that was pretreated with a 2% (v/v) (3-cyanopropyl)-dimethylchlorosilane in acetonitrile by EBS to create a moderately hydrophobic surface. The above KCl buffered electrolyte was used to fill the solution reservoir and the GNM capillary. A voltage was applied across the GNM between two Ag/AgCl electrodes placed inside and outside of the capillary. A lipid bilayer was deposited across the GNM orifice as indicated by a resistance increase from ca. 10 MΩ (associated with the open GNM) to ca. 100 GΩ. A pressure of 20 to 40 mmHg was applied to the inside of the GNM capillary using a syringe, allowing the lipid bilayer to be functional for the protein channel reconstitution. Next, 0.5 µL of α-HL monomer solution at 1 mg/mL was added to the cis side of GNM (a volume of about 150 µL). A voltage of 120 mV (trans vs cis) or higher was applied. The *i-t* traces were filtered typically at 10 kHz and sampled at 50 kHz. Events were extracted using QuB (version 1.5.0.31), and histograms were analyzed by Origin 9.1. Heat plots were plotted using data analysis programs provided by EBS.

#### Single molecule translocations with the CsGg/MinION (ONT platform)

ONT instructions were followed in order to remove air bubbles from the flow cell, add ONT running buffer (RRB), add sample, clean the device after use, and store it. Added samples were either intact oligos or Yenos proprietary osmylated RNA oligos; all the samples were added as is, typically at 1 to 2 µM concentration in ONT running buffer. No library was prepared, and no processing enzyme was added to the sample or buffer, so that all the translocations reported here are unassisted and voltage-driven. Raw data files were acquired in *fast-5* format, visualized directly in MatLab (Mathworks) 2D format. *I*_*r*_, *I*_*o*_, and *τ* values for each event were visually estimated from the grid at apparent accuracy better than the one of the device. Microsoft Excel was used to obtain the histograms as shown in the figures (part D) in the Supplementary Information and plot the normalized counts *vs I*_*r*_*/I*_*o*_ bin in Figure 6. *Fast-5* files were transferred to *txt* format, loaded to QuB open source software (QuB version 1.4.0.1000, the MLab Edition from the Milescu Lab, University of Misssouri and the Sachs Lab at SUNY Buffalo) and concatenated (see Figure 4), as needed.

## Data availability

Some of the data, about 3%, generated during this study are included in this published article (and its Supplementary Information files). All of raw data (each file is about 4GB) may be obtained by request from AK.

## Supplementary Information

see separate file

### Acknowledgements

This work and the authors were supported by NIH/NHGRI grant R43HG010051. AK acknowledges consultation by Dr. Eric Ervin and training by Ryan Dunnam from EBS. AK is grateful to Professor Mark Akeson and Dr. Miten Jain of the UC Santa Cruz Genomics Institute, University of California at Santa Cruz for sharing their software, to visualize raw *i-t* data using MatLab format, with us months before ONT incorporated such capability in their acquisition files. AK acknowledges the skillful assistance of Janette Bernasconi in conducting experiments with the MinION, and of Salma Qureshi in reporting the MinION conductance data. The authors are grateful to Dr. Rehman Ashraf for file formatting and file transferring between MatLab, Microsoft Excel, and QuB open source.

## Author Contributions Statement

K designed and conducted the experiments and supervised data analysis. Both authors reviewed the manuscript.

