## Supplemental Information for "Nanopore device-based fingerprinting of RNA oligos and microRNAs enhanced with an Osmium tag"

### Supplementary Information

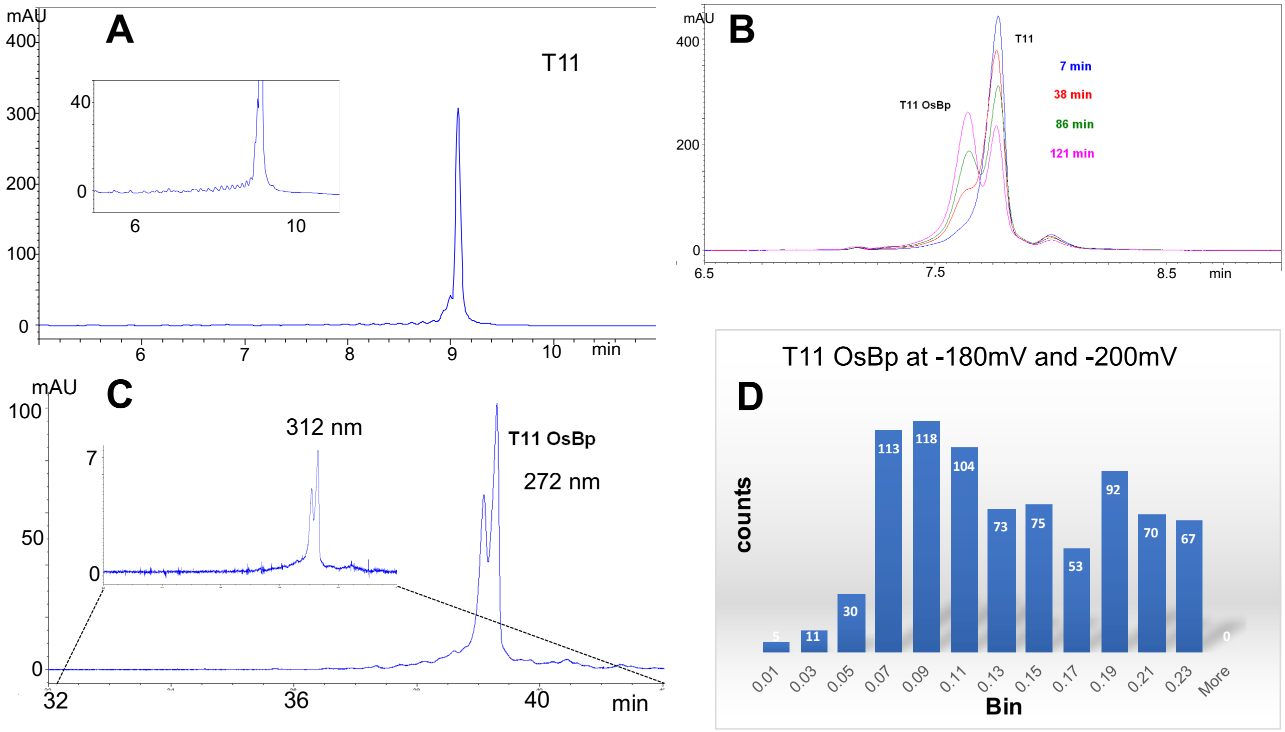


**Figure S1: Analytical profiles of 31nt RNA T11 ($A_{15}CA_{15}$) and osmylated T11 ($A_{15}C(OsBp)A_{15}$).** For details on HPLC, CE and MinION methods and data reporting see Experimental Section.

**A.** IEX pH 8 HPLC profile of 31nt RNA oligo T11 ($A_{15}CA_{15}$) monitored at 260nm. Insert, magnification of the baseline to show one main impurity eluting right before the main peak and many minor peaks attributed to truncated sequences from the synthesis.
**B.** CE profiles monitored at 260nm showing the timely conversion of T11 to the osmylated T11 from the reaction in 5.2mM OsBp. Lower OsBp concentration used here to slow enough the reaction in order to observe the transformation. This CE method using the short capillary does not resolve the two topoisomers, and this is why the product appears as a single peak.
**C.** CE profile of the osmylated T11 ($A_{15}C(OsBp)A_{15}$) monitored at 272nm using the long capillary to enable resolution of the two topoisomers. Insert, same CE profile at 312nm to confirm that both peaks are osmylation products, as intact oligo does not have detectable absorbance at 312nm.
**D.** Fractional residual ion current $I_r/I_o$ histogram with bin=0.02 (811 total count) from the MinION ion channel measurements with $A_{15}C(OsBp)A_{15}$. One experiment conducted at -180mV and data from 5 channels are included; a second experiment conducted at -200mV and data from 2 channels reported here. A plot of normalized counts as a function of $I_r/I_o$ bin is shown in Figure 6A.

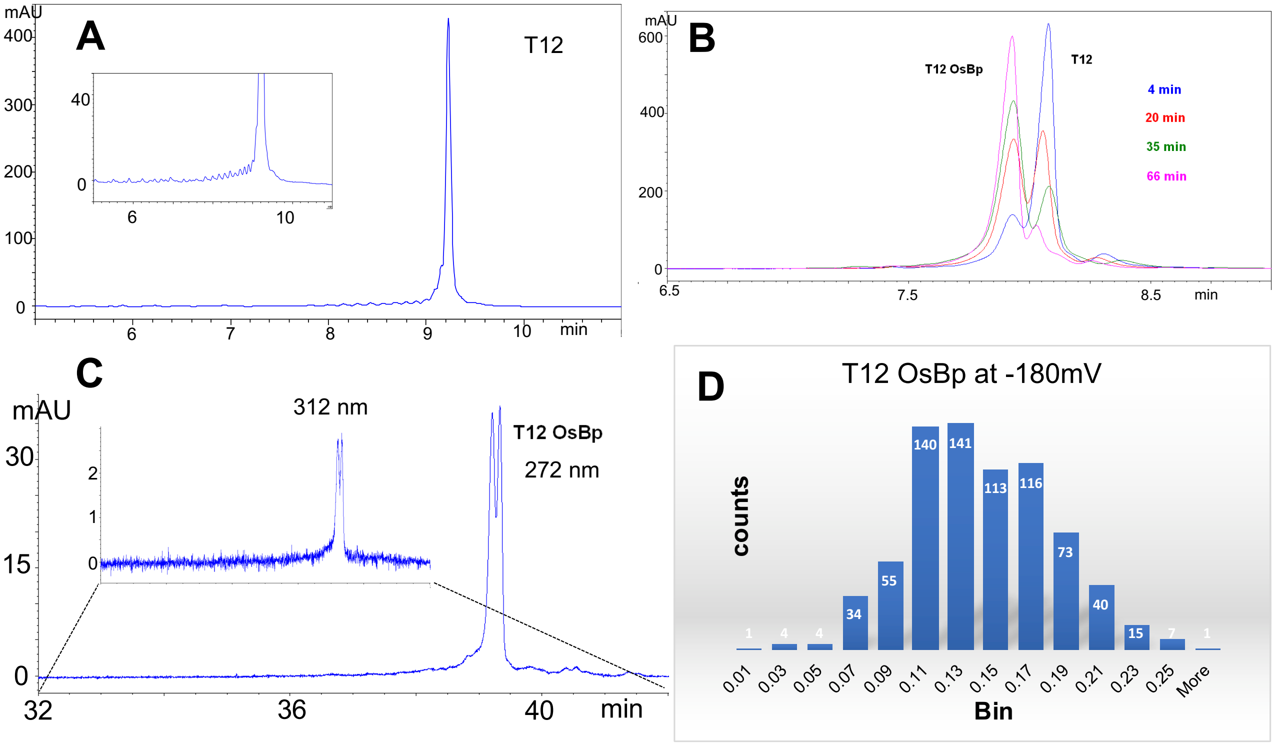


**Figure S2: Analytical profiles of 31nt RNA T12 ($A_{15}UA_{15}$) and osmylated T12 ($A_{15}U(OsBp)A_{15}$).** For details on HPLC, CE and MinION methods and data reporting see Experimental Section.

**A.** IEX pH 8 HPLC profile of 31nt RNA oligo T12 ($A_{15}UA_{15}$) monitored at 260nm. Insert, magnification of the baseline to show minor peaks eluting ahead of the main peak attributed to truncated sequences from the synthesis.
**B.** CE profiles monitored at 260nm showing the timely conversion of T12 to the osmylated T12 from the reaction in 5.2mM OsBp; Lower OsBp concentration used here to slow enough the reaction in order to observe the transformation. This CE method using the short capillary does not resolve the two topoisomers, and this is why the product appears as a single peak.
**C.** CE profile of the osmylated derivative ($A_{15}U(OsBp)A_{15}$) monitored at 272nm using the long capillary to enable resolution of the two topoisomers. Insert, same CE profile at 312nm to confirm that both peaks are osmylation products, as intact oligo does not have detectable absorbance at 312nm.
**D.** Fractional residual ion current $I_r/I_o$ histogram with bin=0.02 (744 total count) from the MinION ion channel measurements with $A_{15}U(OsBp)A_{15}$. One experiment conducted at -180mV and data from 5 channels are reported. A plot of normalized counts as a function of $I_r/I_o$ bin is shown in Figure 6A.

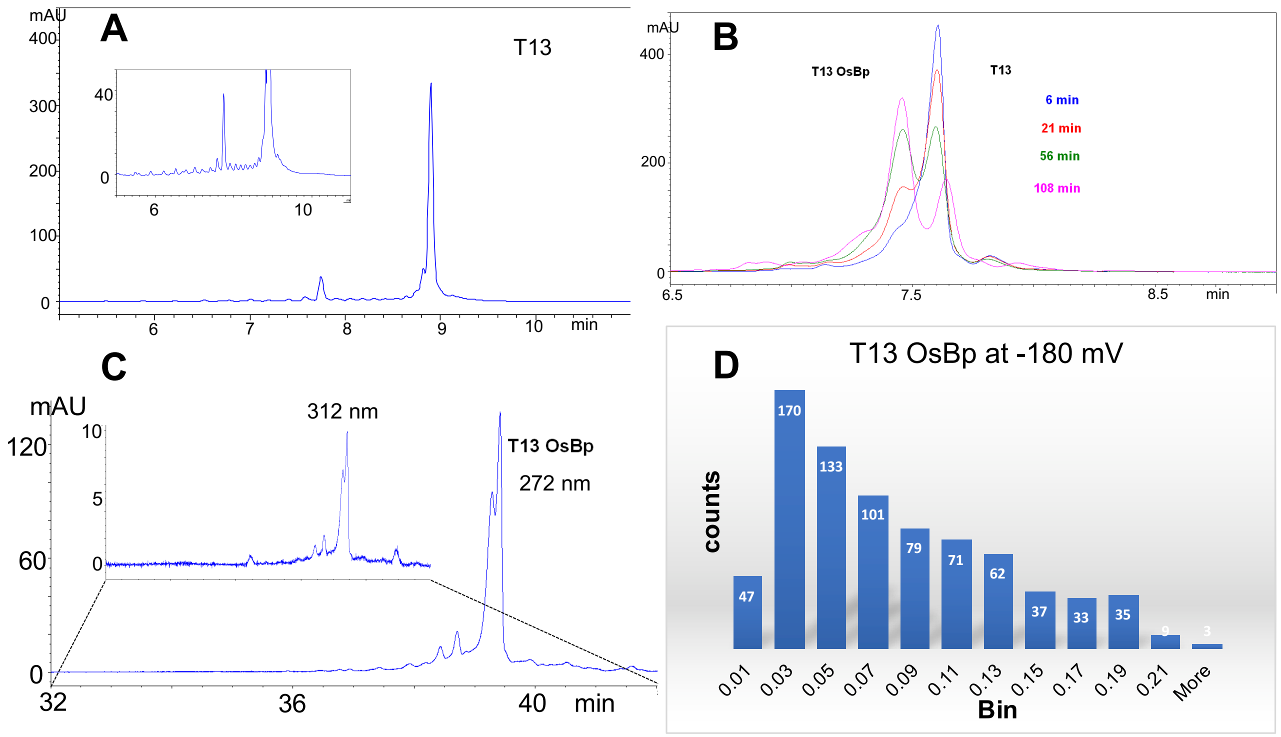


**Figure S3: Analytical profiles of 31nt RNA T13 ($A_{15}$(5Me-C)$A_{15}$) and osmylated T13 ($A_{15}$(5Me-C)(OsBp)$A_{15}$).**

For details on HPLC, CE and MinION methods and data reporting see Experimental Section.

**A.** IEX pH 8 HPLC profile of 31nt RNA oligo T13 ($A_{15}$(5Me-C)$A_{15}$) monitored at 260nm. Insert, magnification of the baseline to show an early eluting impurity as well as minor peaks eluting ahead of the main peak attributed to truncated sequences from the synthesis.

**B.** CE profiles monitored at 260nm showing the timely conversion of T13 to the osmylated T13 from the reaction in 5.2mM OsBp; Lower OsBp concentration used here to slow enough the reaction in order to observe the transformation. This CE method using the short capillary does not resolve the two topoisomers.

**C.** CE profile of the osmylated product ($A_{15}$(5-MeC)(OsBp)$A_{15}$) monitored at 272nm. Insert, same profile at 312nm. The observed two main peaks with no baseline resolution are attributed to the two topoisomers. The early eluting impurity appears to include the pyrimidine, as it exhibits absorbance both at 272nm and at 312nm.

**D.** Fractional residual ion current $I_r/I_o$ histogram with bin=0.02 (780 total count) from the MinION ion channel measurements with $A_{15}$(5-MeC)(OsBp)$A_{15}$. Two experiments conducted at -180mV, and data from 3+2 channels are reported. Visual inspection of the histogram suggests the presence of one major population with $(I_r/I_o)_{max}$ = 0.03. A plot of normalized counts as a function of $I_r/I_o$ bin is shown in Figure 6D.

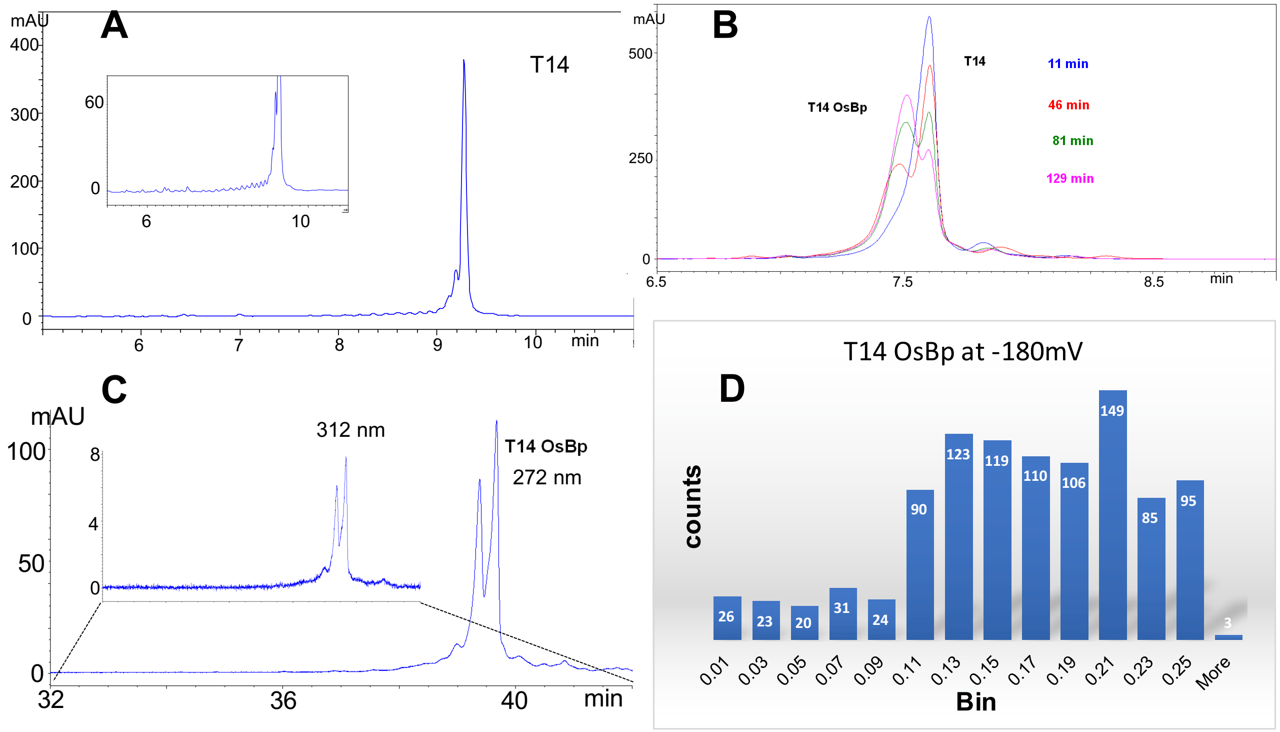


**Figure S4: Analytical profiles of 31nt RNA T14 ($A_{14}GCGA_{14}$) and osmylated T14 ($A_{14}GC(OsBp)GA_{14}$).**
For details on HPLC, CE and MinION methods and data reporting see Experimental Section.

**A.** IEX pH 8 HPLC profile of 31nt RNA oligo T14 ($A_{14}GCGA_{14}$) monitored at 260nm. Insert, magnification of the baseline to show one major impurity eluting right in front of the main peak and minor peaks attributed to truncated sequences from the synthesis.
**B.** CE profiles monitored at 260nm showing the timely conversion of T14 to the osmylated T14 from the reaction in 5.2mM OsBp. Lower OsBp concentration used here to slow enough the reaction in order to observe the transformation. This CE method uses the short capillary and does not resolve the two topoisomers, and this is why the product appears as a single peak.
**C.** CE profile of the osmylated product ($A_{14}GC(OsBp)GA_{14}$) monitored at 272nm using the long capillary for improved resolution. Insert, same profile at 312nm; two main peaks at a ratio of about 1:1 are attributed to the two topoisomers.
**D.** Fractional residual ion current $I_r/I_o$ histogram with bin=0.02 (1004 total count) from the MinION ion channel measurements with $A_{14}GC(OsBp)GA_{14}$. Two experiments both conducted at -180mV, and data were reported from 4 channels each. A plot of normalized counts as a function of $I_r/I_o$ bin is shown in Figure 6B.

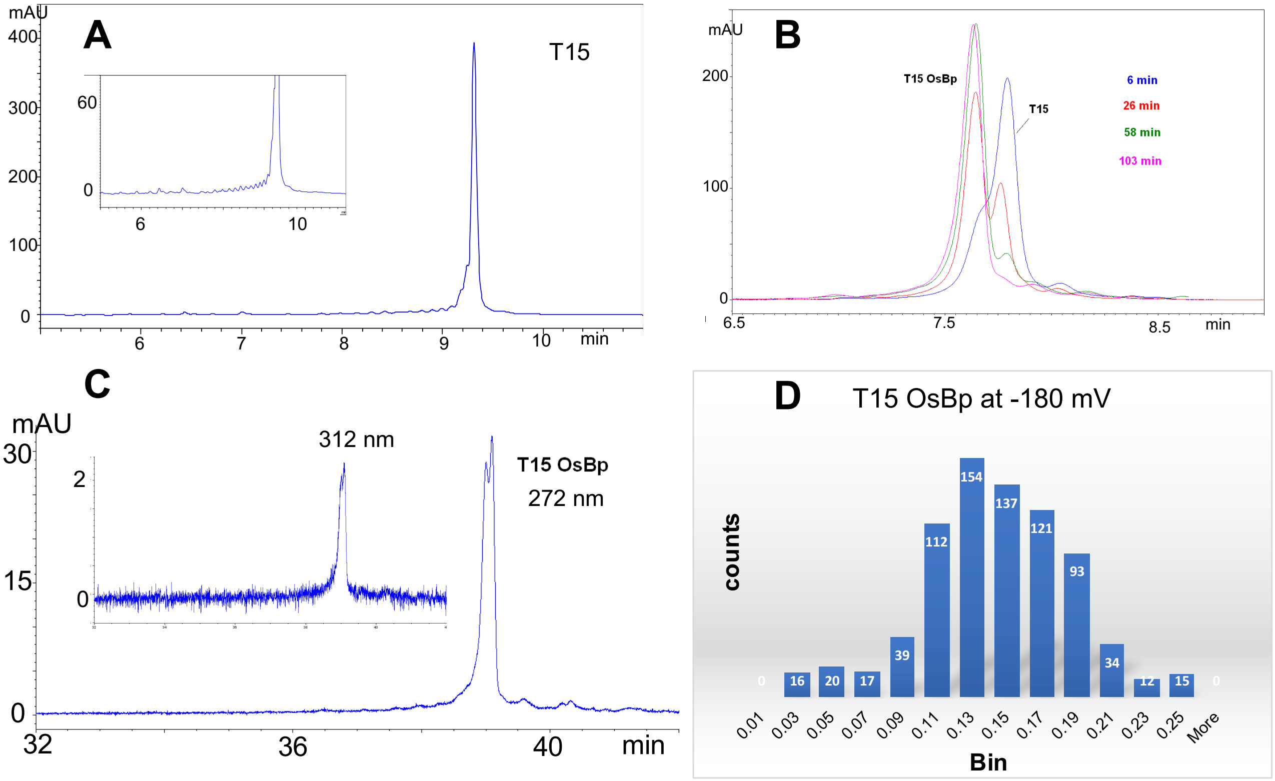


**Figure S5: Analytical profiles of 31nt RNA T15 ($A_{14}GUGA_{14}$) and osmylated T15 ($A_{14}GU(OsBp)GA_{14}$).**
For details on HPLC, CE and MinION methods and data reporting see Experimental Section.

**A.** IEX pH 8 HPLC profile of 31nt RNA oligo T15 ($A_{14}GUGA_{14}$) monitored at 260nm. Insert, magnification of the baseline to show minor peaks eluting ahead of the main peak attributed to truncated sequences from the synthesis.
**B.** CE profiles monitored at 260nm showing the timely conversion of T15 to the osmylated T15 from the reaction in 5.2mM OsBp. Lower OsBp concentration used here to slow enough the reaction in order to observe the transformation. This CE method uses the short capillary and does not resolve the two topoisomers.
**C.** CE profile of the osmylated derivative ($A_{14}GU(OsBp)GA_{14}$) monitored at 272nm using the long capillary for improved resolution. Insert, same profile at 312nm; two peaks at a ratio of about 1:1 are attributed to the two topoisomers.
**D.** Fractional residual ion current $I_r/I_o$ histogram with bin=0.02 (770 total count) from the MinION ion channel measurements with $A_{14}GU(OsBp)GA_{14}$. Two experiments both conducted at -180mV, and data were reported from 5 + 2 channels. A plot of normalized counts as a function of $I_r/I_o$ bin is shown in Figure 6C.

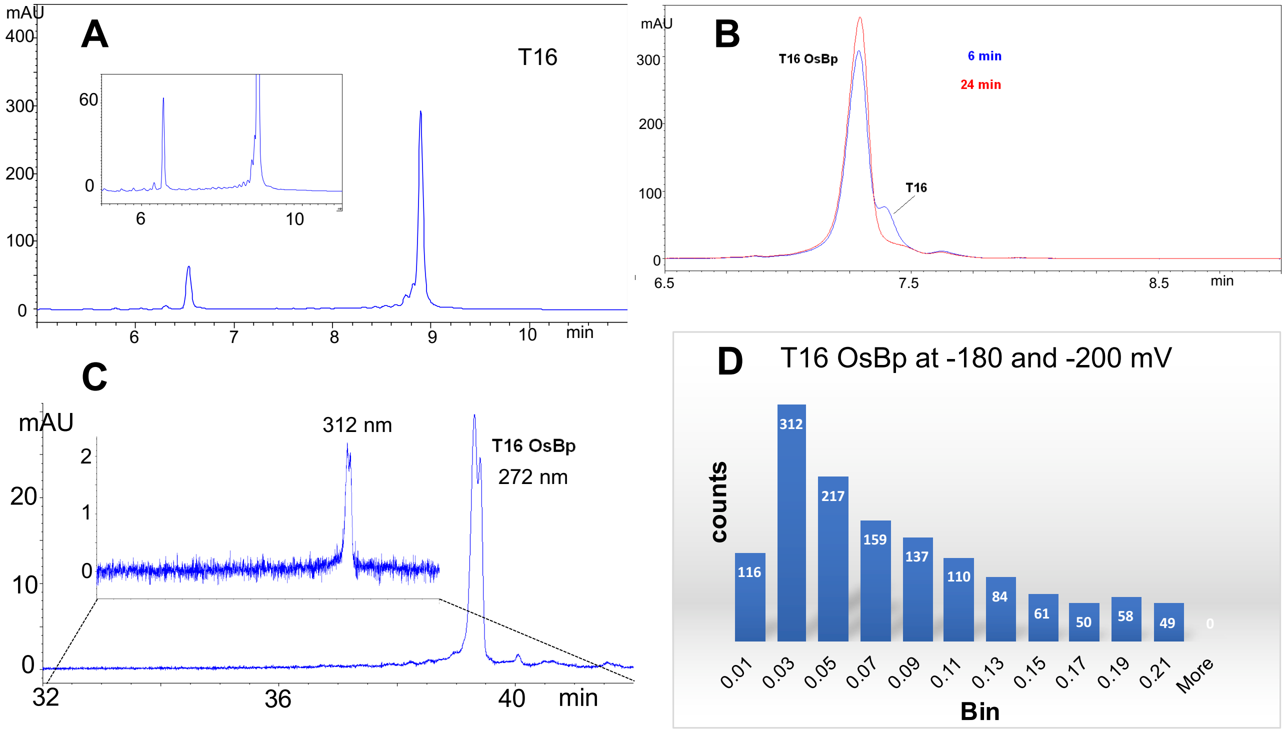


**Figure S6: Analytical profiles of 31nt RNA T16 ($A_{15}$(5-MeU)$A_{15}$) and osmylated T16 ($A_{15}$(5-MeU)(OsBp)$A_{15}$).**

For details on HPLC, CE and MinION methods and data reporting see Experimental Section.

**A.** IEX pH 8 HPLC profile of 31nt RNA oligo T16 ($A_{15}$(5-MeU)$A_{15}$) monitored at 260nm. Insert, magnification of the baseline to show an early eluting impurity as well as minor peaks eluting ahead of the main peak attributed to truncated sequences from the synthesis. The early eluting impurity appears to include no pyrimidine, as it exhibits no absorbance at 312nm (not shown).

**B.** CE profiles monitored at 260nm showing the conversion of T16 to the osmylated T16 from the reaction in 5.2mM OsBp; only two profiles are shown as the reaction is very fast and close to complete in 6 minutes.

**C.** CE profile of the product ($A_{15}$(5-MeU)(OsBp)$A_{15}$) monitored at 272nm using the long capillary for improved resolution. Insert, same profile at 312nm; two main peaks at a ratio of about 1:1 are attributed to the two topoisomers.

**D.** Fractional residual ion current $I_r/I_o$ histogram with bin=0.02 (1353 total count) from the MinION ion channel measurements with $A_{15}$(5-MeU)(OsBp)$A_{15}$. Three experiments conducted at -180mV, and one experiment at -200mV; data are reported from 3+3+1+1 channels, respectively. Visual inspection of the histogram suggests the presence of one major population with $(I_r/I_o)_{max}$ = 0.03. A plot of normalized counts as a function of $I_r/I_o$ bin is shown in Figure 6D.

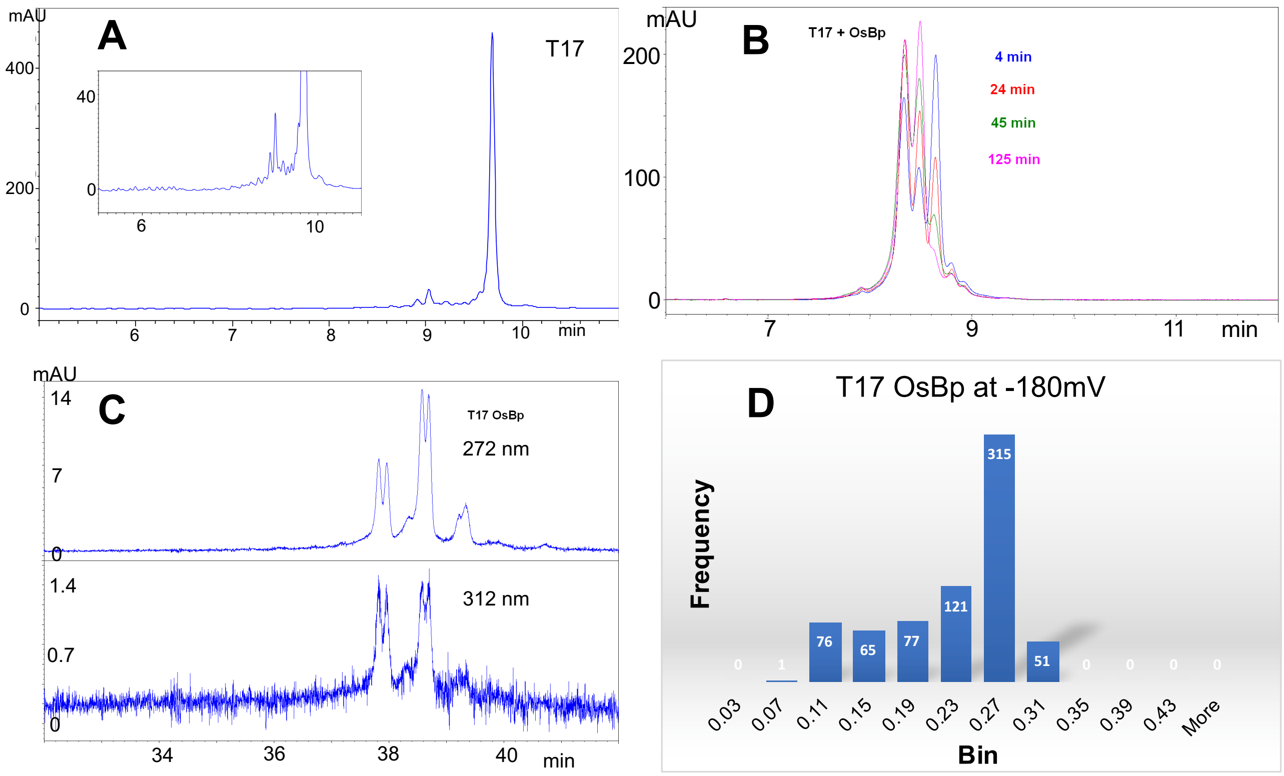


**Figure S7: Analytical profiles of 31nt RNA T17 ($A_{15}$4-SU$A_{15}$) and osmylated T17 ($A_{15}$4-SU(OsBp)$A_{15}$).**
For details on HPLC, CE and MinION methods and data reporting see Experimental Section.

**A.** IEX pH 8 HPLC profile of 31nt RNA oligo T17 ($A_{15}$4-SU$A_{15}$) monitored at 260nm. Insert, magnification of the baseline to show an impurity and minor peaks eluting ahead of the main peak attributed to truncated sequences from the synthesis. T17 measures R(312/272) =0.05, because the 4S-U moiety has $\lambda_{max}$ at 331 nm; all the other RNAs tested measure R(312/272) =0.01.
**B.** CE profiles monitored at 260nm showing the timely conversion of T17 to the osmylated T17 from the reaction in 6.3 mM OsBp; Lower OsBp concentration used here to slow enough the reaction in order to observe the transformation. The peak at the far right corresponds to intact T17 and the two ealier migrating peaks correspond to two osmylated products (see text). Impurities in the intact material do not significantly contribute to this transformation. This CE method uses the short capillary and does not resolve topoisomers.
**C.** CE profiles of the osmylated products ($A_{15}$4-SU(OsBp)$A_{15}$): Top profile monitored at 272nm using the long capillary for improved resolution; bottom, at 312nm. Two products are confirmed with two peaks each at about 1:1 ratio attributed to two topoisomers per product. The earlier migrating product exhibits relatively more absorbance at 312nm compared to the later migrating product.
**D.** Fractional residual ion current $I_r/I_o$ histogram with bin=0.04 (706 total count) from the MinION ion channel measurements with $A_{15}$4-SU(OsBp)$A_{15}$. One experiment conducted at -180mV, and data from 2 channels are reported. Visual inspection of the histogram suggests the presence of a population with $(I_r/I_o)_{max}$ = 0.27. A plot of normalized counts as a function of $I_r/I_o$ bin is shown in Figure 6E.

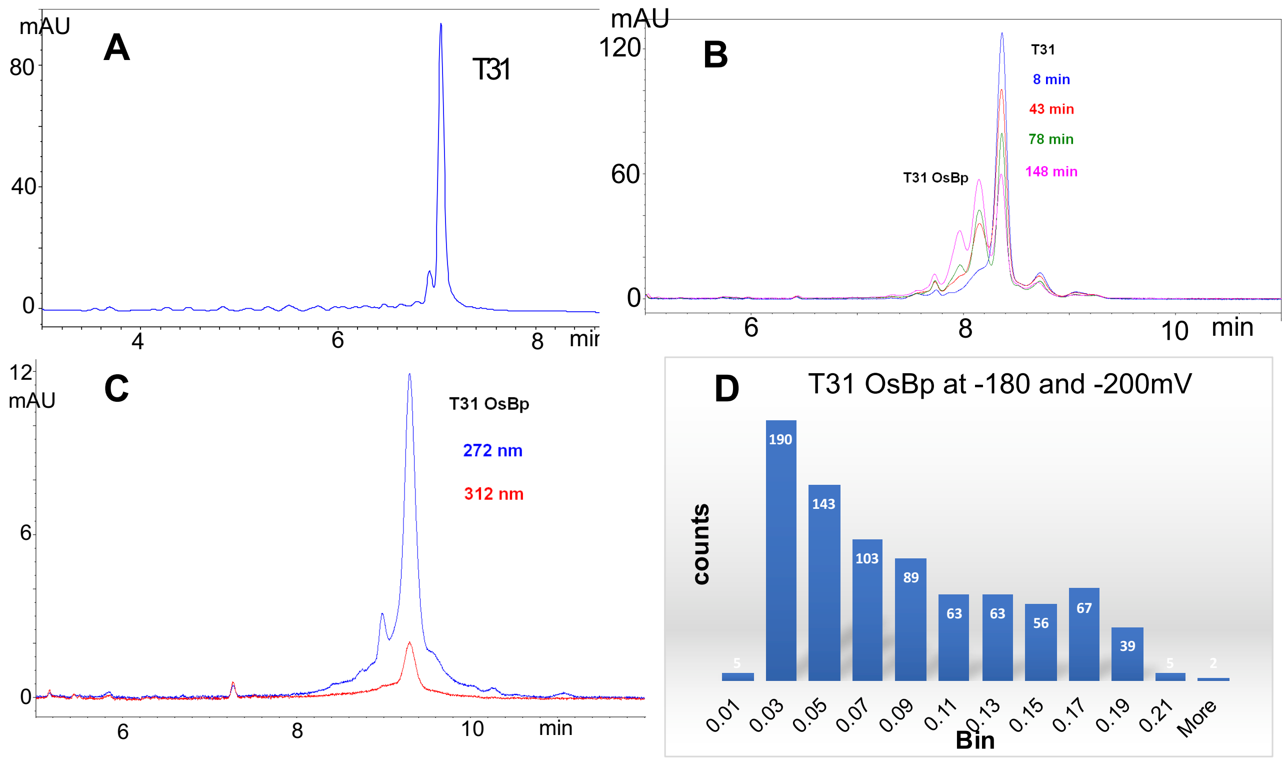


**Figure S8: Analytical profiles of 22nt RNA T31 ($A_{10}CCA_{10}$) and osmylated T31 ($A_{10}C^*C^*A_{10}$); *stands for OsBp.**

For details on HPLC, CE and MinION methods and data reporting see Experimental Section.

**A.** IEX pH 8 HPLC profile of 31nt RNA oligo T31 ($A_{10}CCA_{10}$) monitored at 260nm. Insert, magnification of the baseline to show an impurity as well as minor peaks eluting ahead of the main peak attributed to truncated sequences from the synthesis.

**B.** CE profiles monitored at 260nm showing the timely conversion of T31 to the osmylated T31 from the reaction in 6.3 mM OsBp. CE resolves the singly, forming first (later migration time (m.t.)), from the doubly osmylated product (earlier m.t.), forming second. The more OsBp moieties on a molecule the earlier the corresponding m.t. Reaction shown does not go to completion.

**C.** CE profile of the fully osmylated product ($A_{10}C^*C^*A_{10}$) monitored at 272nm with the short capillary. Insert, same profile at 312nm; quantitation in Table 1. For discussion on topoisomers see Figure 3B and text.

**D.** Fractional residual ion current $I_r/I_o$ histogram with bin=0.02 (825 total count) from the MinION ion channel measurements with $A_{10}C^*C^*A_{10}$. Two experiments were conducted, one at -180mV and the other at -200mV and 4+2 channels are reported. A plot of normalized counts as a function of $I_r/I_o$ bin is shown in Figure 6F.

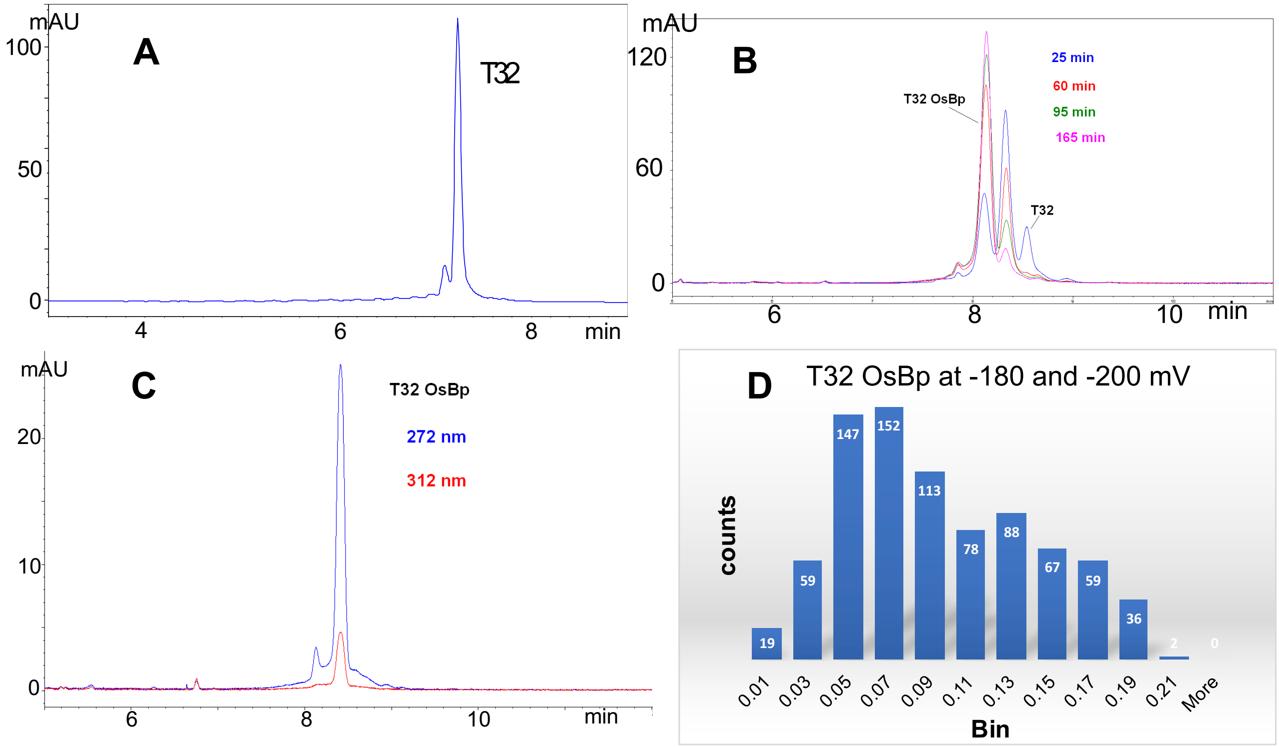


**Figure S9: Analytical profiles of 22nt RNA T32 ($A_{10}UUA_{10}$) and osmylated T32 ($A_{10}U^*U^*A_{10}$); * stands for OsBp.**
For details on HPLC, CE and MinION methods and data reporting see Experimental Section.

**A.** IEX pH 8 HPLC profile of 32nt RNA oligo T32 ($A_{10}UUA_{10}$) monitored at 260nm; this HPLC method resolves well T32 from T31 (not shown).
**B.** CE profiles monitored at 260nm showing the timely conversion of T32 to the osmylated T32 from the reaction in 6.3 mM OsBp. CE resolves the singly, forming first (later migration time (m.t.)), from the doubly osmylated product (earlier m.t.), forming second. The more OsBp moieties on a molecule the earlier the corresponding m.t. Reaction shown does not go to completion, but it is quite a bit faster compared to the one with T31 (Figure S8.B), as U osmylates 4.7-times faster compared to C.
**C.** CE profile of the fully osmylated product ($A_{10}U^*U^*A_{10}$) monitored at 272nm with the short capillary. Insert, same profile at 312nm; quantitation in Table 1. For discussion on topoisomers see Figure 3B and text.
**D.** Fractional residual ion current $I_r/I_o$ histogram with bin=0.02 (820 total count) from the MinION ion channel measurements with $A_{10}U^*U^*A_{10}$. Two experiments were conducted, one at -180mV and the other at -200mV and 9+1 channels are reported. A plot of normalized counts as a function of $I_r/I_o$ bin is shown in Figure 6F.

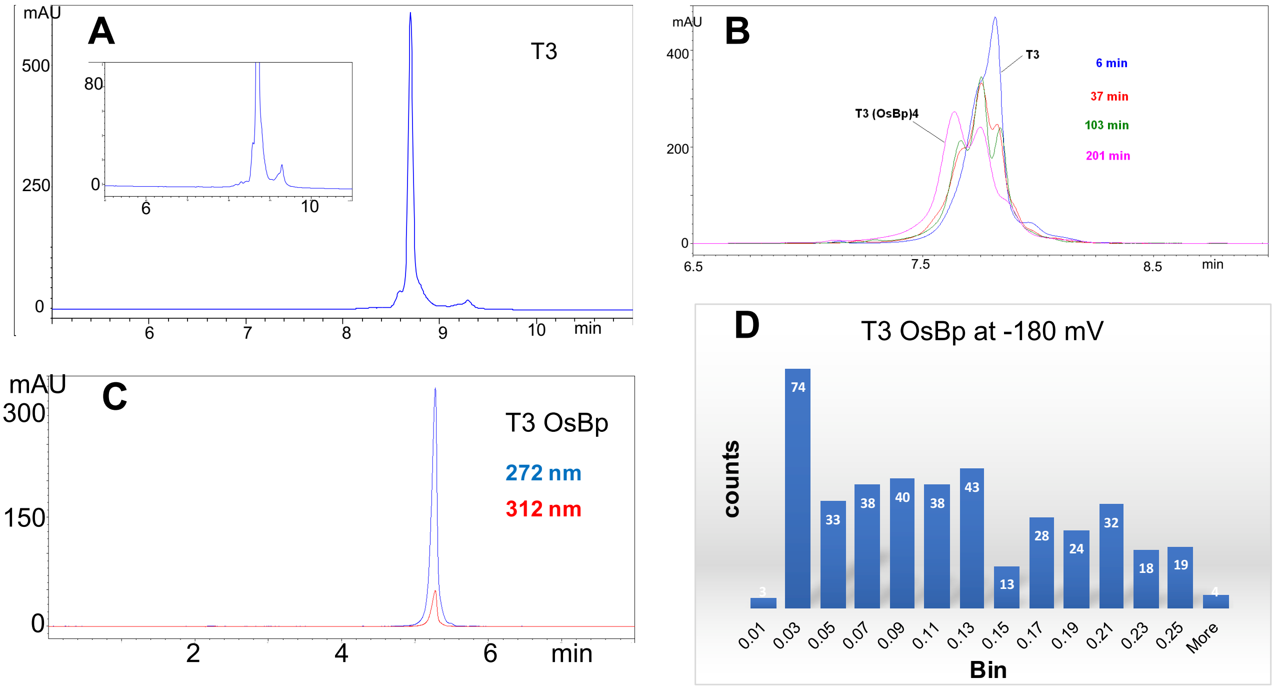


**Figure S10: Analytical profiles of 53nt RNA T3 ($A_{15}UUA_{19}UUA_{15}$) and osmylated T3 ($A_{15}U^*U^*A_{19}U^*U^*A_{15}$); * stands for OsBp.**
For details on HPLC, CE and MinION methods and data reporting see Experimental Section.

**A.** IEX pH 12 HPLC profile of 53nt RNA oligo T3 ($A_{15}UUA_{19}UUA_{15}$) obtained at 260nm. Insert, magnification of the baseline to show the impurity profile; intact oligo measures R(312/272) =0.01.
**B.** CE profiles monitored at 260nm showing the timely conversion of T3 to the osmylated T3 from the reaction in 5.2 mM OsBp. Reaction is relatively slow at these conditions and partially osmylated products can be observed to form and react further; more osmylation leads to peaks migrating at earlier times. Fully osmylated T3 includes 4 OsBp moieties. Reaction does not go to completion under these conditions. Manufacturing was conducted by 3h incubation in 10.5 mM OsBp (see C.).
**C.** CE profile of the fully osmylated product ($A_{15}U^*U^*A_{19}U^*U^*A_{15}$) monitored at 272nm (blue) with the short capillary and same profile at 312nm (red); quantitation in Table 1. For discussion on topoisomers see Figure 3B and text.
**D.** Fractional residual ion current $I_r/I_o$ histogram with bin=0.02 (407 total count) from the MinION ion channel measurements with $A_{15}U^*U^*A_{19}U^*U^*A_{15}$. One experiment was conducted at -180mV and two channels are reported. More than one $(I_r/I_o)_{max}$ are consistent with the topoisomerism discussed in the text. A plot of normalized counts as a function of $I_r/I_o$ bin is shown in Figure 6G.

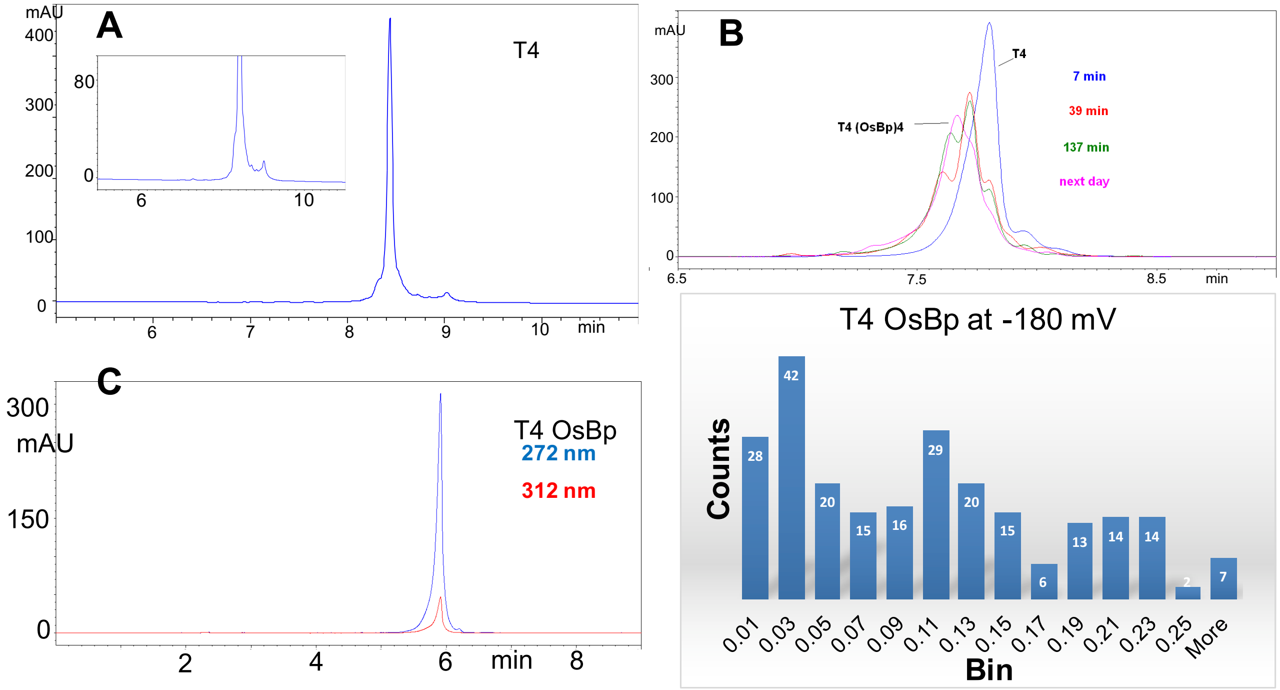


**Figure S11: Analytical profiles of 53nt RNA T4 ($A_{15}CCA_{19}UUA_{15}$) and osmylated T4 ($A_{15}C^*C^*A_{19}U^*U^*A_{15}$); * stands for OsBp.**
For details on HPLC, CE and MinION methods and data reporting see Experimental Section.

**A.** IEX pH 12 HPLC profile of 53nt RNA oligo T4 ($A_{15}CCA_{19}UUA_{15}$) obtained at 260nm. Insert, magnification of the baseline to show the impurity profile; intact oligo measures R(312/272) =0.01.
**B.** CE profiles monitored at 260nm showing the timely conversion of T4 to the osmylated T4 from the reaction in 5.2 mM OsBp. Reaction is relatively slow at these conditions and partially osmylated products can be observed to form and react further; more osmylation leads to peaks migrating at earlier times. Fully osmylated T4 includes 4 OsBp moieties. Reaction does not go to completion under these conditions. Manufacturing was conducted by 3h incubation in 10.5 mM OsBp (see C.).
**C.** CE profile of the fully osmylated product ($A_{15}C^*C^*A_{19}U^*U^*A_{15}$) monitored at 272nm with the short capillary (blue) and same profile at 312nm (red); For discussion on topoisomers see Figure 3B and text.
**D.** Fractional residual ion current $I_r/I_o$ histogram with bin=0.02 (241 total count) from the MinION ion channel measurements with $A_{15}C^*C^*A_{19}U^*U^*A_{15}$. One experiment was conducted at -180mV and 10 channels are reported. Material was used at about 0.1μM concentration (1/10 dilution over the typical concentration used) and the effect was visible in obtaining very few translocations per unit time. More than one $(I_r/I_o)_{max}$ are consistent with the topoisomerism discussed in the text.

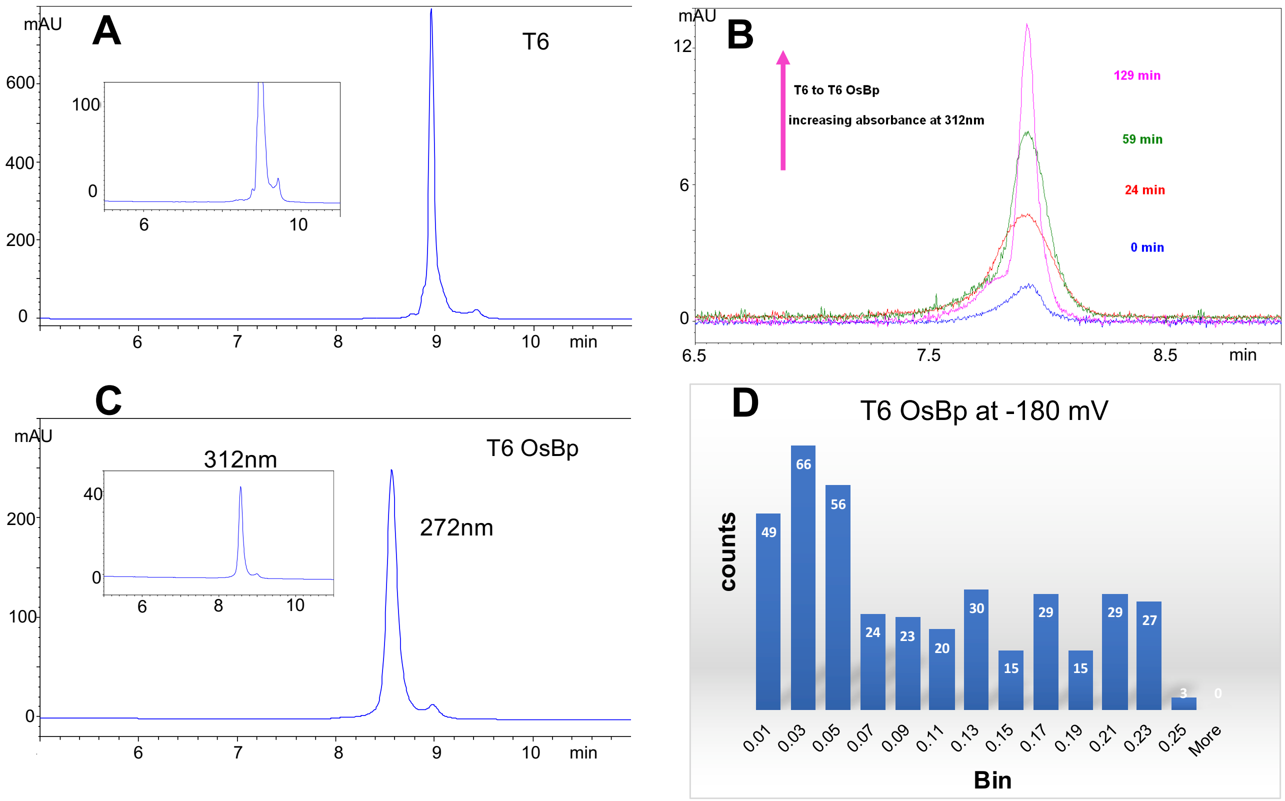


**Figure S12: Analytical profiles of 74nt RNA T6 ($A_{15}UUA_{19}UUA_{19}UUA_{15}$) and osmylated T6 ($A_{15}U*U*A_{19}U*U*A_{19}U*U*A_{15}$); * stands for OsBp.**

For details on HPLC and CE Methods and data reporting see Experimental Section.

**A.** IEX pH 12 HPLC profile of 74nt RNA oligo T6 ($A_{15}UUA_{19}UUA_{19}UUA_{15}$) obtained at 260nm. Insert, magnification of the baseline to show the impurity profile; intact oligo measures R(312/272) =0.01.

**B.** UV-Vis Absorbance of peak monitored at 312nm by CE showing the timely conversion of T6 to the osmylated T6 from the reaction in 5.2 mM OsBp. Too many partially osmylated products are formed and can't be resolved by CE, but the observed peak moves to earlier m.t. with time, not replicated here. Fully osmylated T6 includes 6 OsBp moieties. Reaction does not go to completion under these conditions. Manufacturing was conducted by 3h incubation in 10.5 mM OsBp (see C.).

**C.** CE profile of the fully osmylated product ($A_{15}U*U*A_{19}U*U*A_{19}U*U*A_{15}$) monitored at 272nm with the short capillary. Insert, same profile at 312nm; quantitation in Table 1. For discussion on topoisomers see Figure 3B and text.

**D.** Fractional residual ion current $I_r/I_o$ histogram with bin=0.02 (386 total count) from the MinION ion channel measurements with $A_{15}U*U*A_{19}U*U*A_{19}U*U*A_{15}$. One experiment was conducted at -180mV and 3 channels are reported. A plot of normalized counts as a function of $I_r/I_o$ bin is shown in Figure 6G.

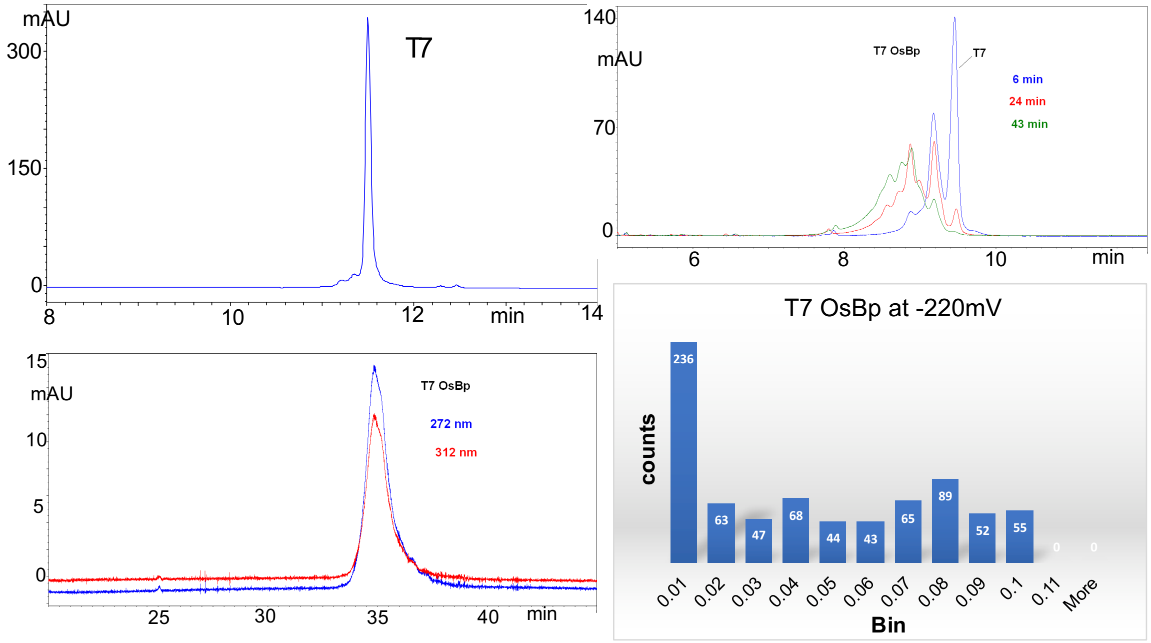


**Figure S13: Analytical profiles of 32nt RNA T7(13Py) ($(AG)_3C_4(AG)_3C_4(AG)_3CCUUCA$) and osmylated T7(13Py) with 13 pyrimidines practically 100% osmylated.**

For details on HPLC, CE, and MinION methods and data reporting see Experimental Section.

**A.** IEX pH 12 HPLC profile of 32nt RNA oligo T7(13Py) monitored at 260nm; this HPLC method resolves well T7 from T8 (not shown).

**B.** CE profiles monitored at 260nm showing the timely conversion of T8 to the osmylated T8 from the reaction in 6.3 mM OsBp. CE resolves the singly, forming first (later migration time (m.t.)), from the later forming more osmylated products (earlier m.t.). The more OsBp moieties on a molecule the earlier the corresponding m.t. Reaction shown does not go to completion. Manufacturing 3h in 12.6 mM OsBp.

**C.** CE profiles of the fully osmylated T7 with 13 OsBp moieties monitored at 272nm with the long capillary at 272nm (blue trace) and at 312nm (red trace); quantitation in Table 1. Resolution can't be achieved due to the multitude of topoisomers.

**D.** Fractional residual ion current $I_r/I_o$ histogram with bin=0.01 (762 total count) from the MinION ion channel measurements with osmylated T7. One experiments was conducted at -220mV and seven channels are reported. A plot of normalized counts as a function of $I_r/I_o$ bin is shown in Figure 6H.

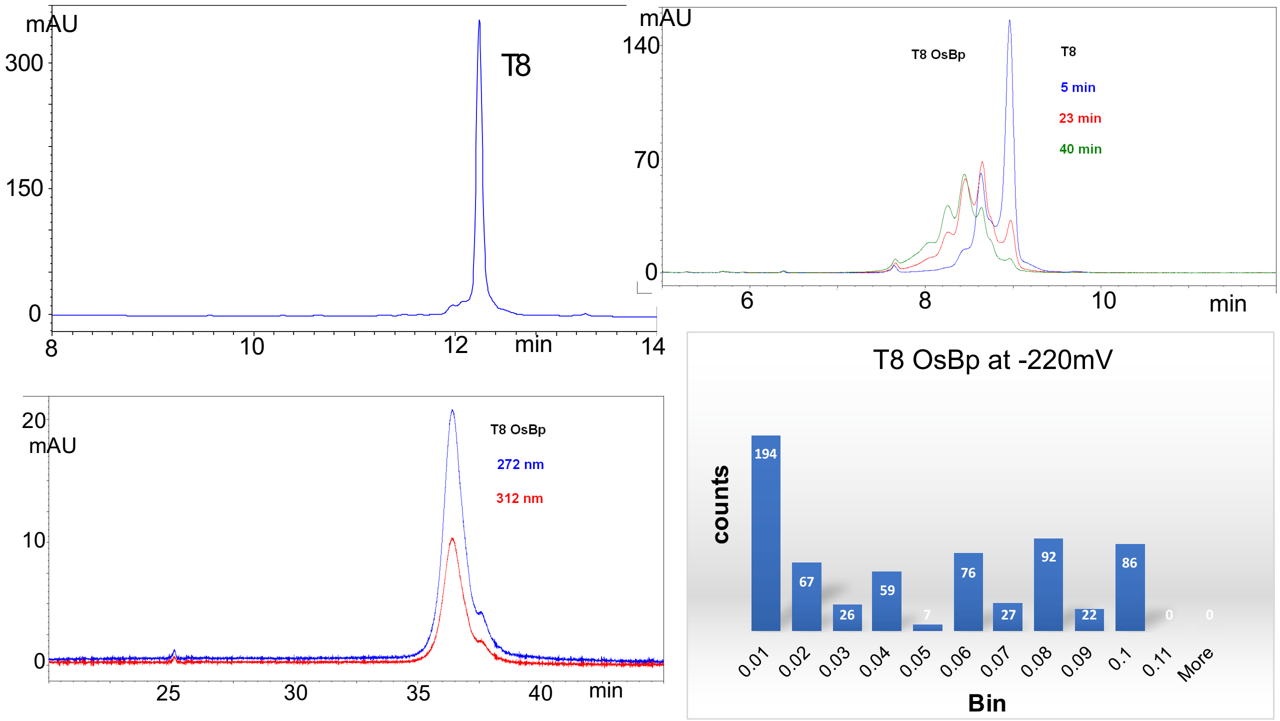


**Figure S14: Analytical profiles of 32nt RNA T8(9Py) ($(AG)_4C_2(AG)_4C_2(AG)_3$CCUUCA) and osmylated T8(9Py) with 9 pyrimidines practically 100% osmylated.**

For details on HPLC, CE, and MinION methods and data reporting see Experimental Section.

**A.** IEX pH 12 HPLC profile of 32nt RNA oligo T8(9Py) monitored at 260nm; this HPLC method resolves well T7 from T8 (not shown).

**B.** CE profiles monitored at 260nm showing the timely conversion of T8 to the osmylated T8 from the reaction in 6.3 mM OsBp. CE resolves the singly, forming first (later migration time (m.t.)), from the later forming more osmylated products (earlier m.t.). The more OsBp moieties on a molecule the earlier the corresponding m.t. Reaction shown does not go to completion. Manufacturing 3h in 12.6 mM OsBp.

**C.** CE profile of the fully osmylated T8 with 9 OsBp moieties monitored at 272nm (blue trace) and at 312nm (red trace) with the long capillary; quantitation in Table 1. Resolution can't be achieved due to the multitude of topoisomers.

**D.** Fractional residual ion current $I_r/I_o$ histogram with bin=0.01 (656 total count) from the MinION ion channel measurements with osmylated T8. One experiment was conducted at -220mV and ten channels are reported. A plot of normalized counts as a function of $I_r/I_o$ bin is shown in Figure 6H.

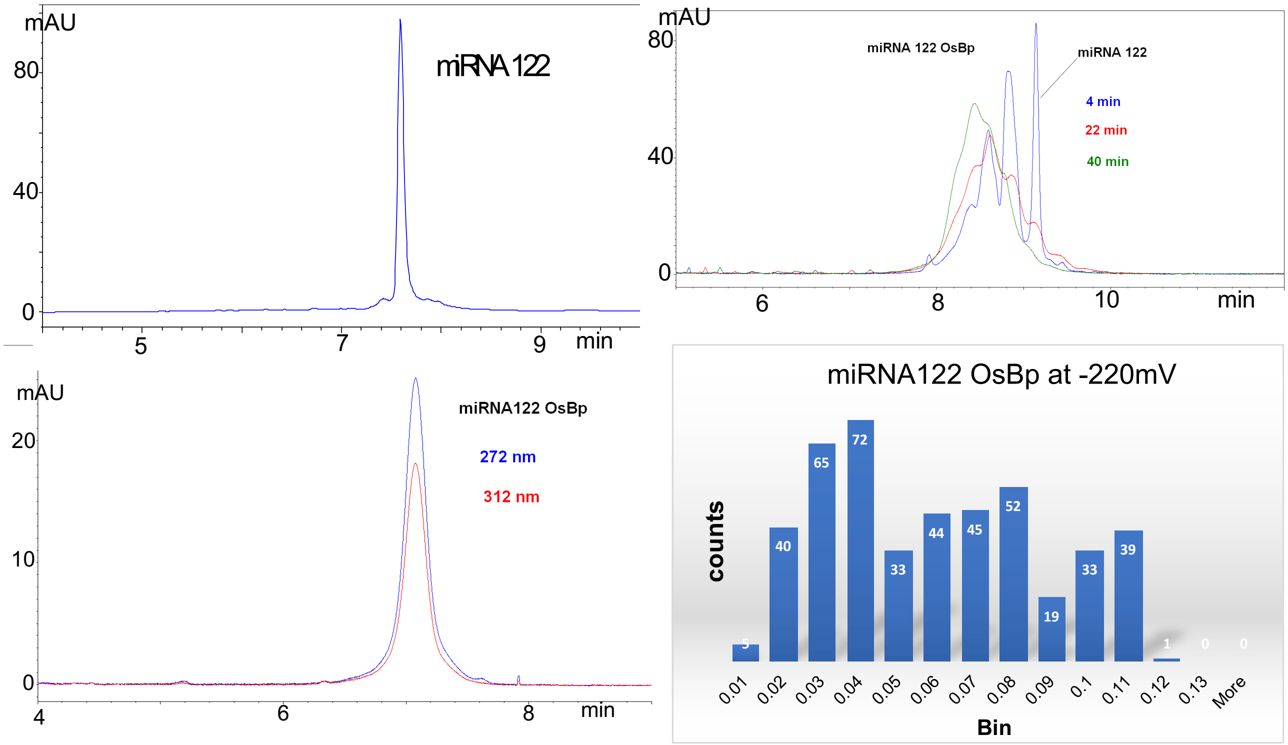


**Figure S15: Analytical profiles of 22nt miRNA 122 (sequence in Table 1) and osmylated miRNA 122 with 9 pyrimidines practically 100% osmylated.**
For details on HPLC, CE, and MinION methods and data reporting see Experimental Section.

**A.** IEX pH 8 HPLC profile of 22nt miRNA 122 monitored at 260nm; this HPLC method resolves miRNA 122 from miRNA 140 (not shown).
**B.** CE profiles monitored at 260nm showing the timely conversion of miRNA 122 to the osmylated products from the reaction in 6.3 mM OsBp. CE resolves the singly, forming first (later migration time (m.t.)), from the later forming more osmylated products (earlier m.t.). The more OsBp moieties on a molecule the earlier the corresponding m.t. Reaction shown here does not go to completion. Manufacturing 3h in 12.6 mM OsBp.
**C.** CE profile of the fully osmylated miRNA 122 with 9 OsBp moieties monitored at 272nm (blue) and at 312nm (red) with the short capillary; quantitation in Table 1.
**D.** Fractional residual ion current $I_r/I_o$ histogram with bin=0.01 (448 total count) from the MinION ion channel measurements with osmylated miRNA 122. One experiment was conducted at -220mV and 3 channels are reported. A plot of normalized counts as a function of $I_r/I_o$ bin is shown in Figure 6I.

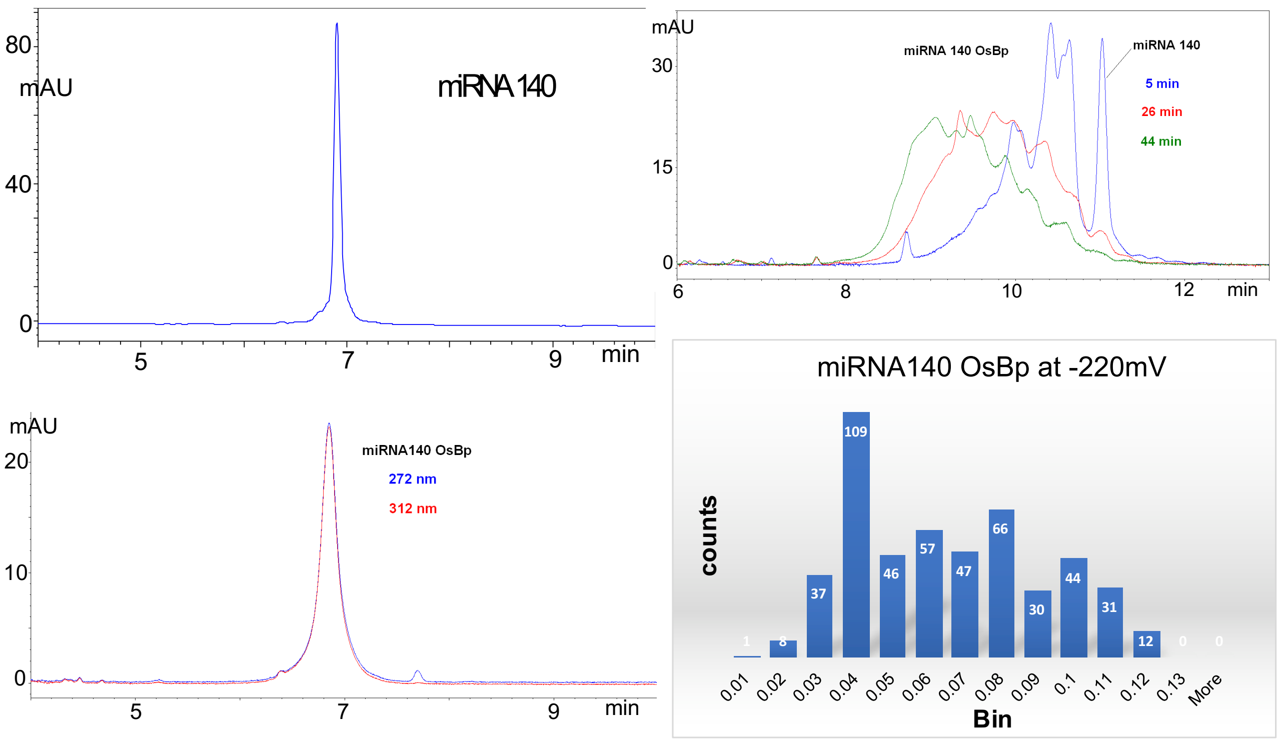


**Figure S16: Analytical profiles of 22nt miRNA 140 (sequence in Table 1) and osmylated miRNA 140 with 12 pyrimidines practically 100% osmylated.**
For details on HPLC, CE, and MinION methods and data reporting see Experimental Section.

**A.** IEX pH 8 HPLC profile of 32nt miRNA 140 monitored at 260nm.
**B.** CE profiles monitored at 260nm showing the timely conversion of miRNA 140 to the osmylated product from the reaction in 6.3 mM OsBp. CE resolves the singly, forming first (later migration time (m.t.)), from the later forming more osmylated products (earlier m.t.). The more OsBp moieties on a molecule the earlier the corresponding m.t. Reaction shown here does not go to completion. Manufacturing 3h in 12.6 mM OsBp.
**C.** CE profile of the fully osmylated miRNA 140 with 12 OsBp moieties monitored at 272nm (blue) and at 312nm (re) with the short capillary, practically overlapping; R(312/272)≈1, see Table 1.
**D.** Fractional residual ion current $I_r/I_o$ histogram with bin=0.01 (488 total count) from the MinION ion channel measurements with osmylated miRNA 140. One experiment was conducted at -220mV and 3 channels are reported. A plot of normalized counts as a function of $I_r/I_o$ bin is shown in Figure 6I.
References for function see:

Zhao, Z., Liu, W. & Li, J. miR-140-5p inhibits cell proliferation and invasion in colorectal carcinoma by targeting SOX4. *Oncol Lett.* **17**(2), 2215-2220, https://doi.org/10.3892/ol.2018.9834 (2019).
.Lu, Y. et al. MicroRNA-140-5p inhibits invasion and angiogenesis through targeting VEGF-A in breast cancer. *Cancer Gene Ther.* **24**(9), 386-392, https://doi.org/10.1038/cgt.2017.30 (2017).
Si, H. B. et al. Intra-articular injection of microRNA-140 (miRNA-140) alleviates osteoarthritis (OA) progression by modulating extracellular matrix (ECM) homeostasis in rats. *Osteoarthritis Cartilage*. **25**(10), 1698-1707, https://doi.org/10.1016/j.joca.2017.06.002. (2017)
Liao, W. et al. MicroRNA-140-5p attenuated oxidative stress in Cisplatin induced acute kidney injury by activating Nrf2/ARE pathway through a Keap1-independent mechanism. *Exp Cell Res.* **360**(2), 292-302, https://doi.org/10.1016/j.yexcr.2017.09.019 (2017).

**Table S1: Pseudo-first order rates of osmylation (1/min) for 20nt deoxyoligos**

| [OsBp], mM[a] | $[OsBp]^2$ | $A_{10}CA_9$, Rate, k (1/min) | $A_{10}dUA_9$, Rate, k (1/min) |
|---|---|---|---|
| 1.575 | 2.48 | 0.00134 | ND |
| 3.15 | 9.92 | 0.0107 | 0.021 |
| 6.3 | 39.69 | 0.023 | 0.084 |
| 9.45 | 89.30 | 0.049 | 0.265[b] |
| 12.6 | 158.76 | 0.085 | ND |

ND stands for not determined

[a] OsBp stock solution prepared from an equimolar mixture of $OsO_4$ and 2,2'-bipyridine; OsBp concentration, nominal [OsBp], with [OsBp]=[$OsO_4$]=[2,2'-bipyridine].

[b] Relatively fast reaction, rate obtained from two data points, 0 and 4 min; accuracy at ± 15%.

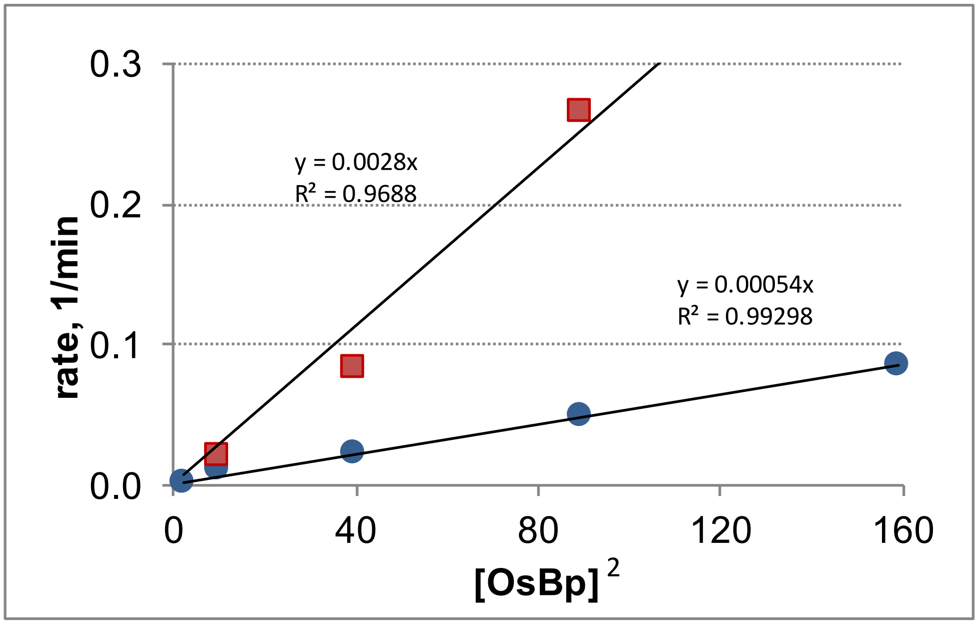


**Figure S17**: **Data from Table S1** (see above). Linear extrapolations go via 0,0 intercept consistent with no reaction in the absence of OsBp, i.e. at [OsBp]=0. Deoxyoligo $A_{10}dUA_9$, (red squares) and deoxyoligo $A_{10}CA_9$, (blue circles), consistent with the observed faster osmylation rate with dU *vs* dC and U *vs* C. The linear square dependence of the osmylation pseudo-first order rate on nominal OsBp concentration suggests that preassociation or complexation of $OsO_4$ with 2,2'-bipyridine is small under these conditions (see text). It is presumed that the same square dependence on [OsBp] applies to the osmylation of RNA oligos, as seen here with deoxyoligos.

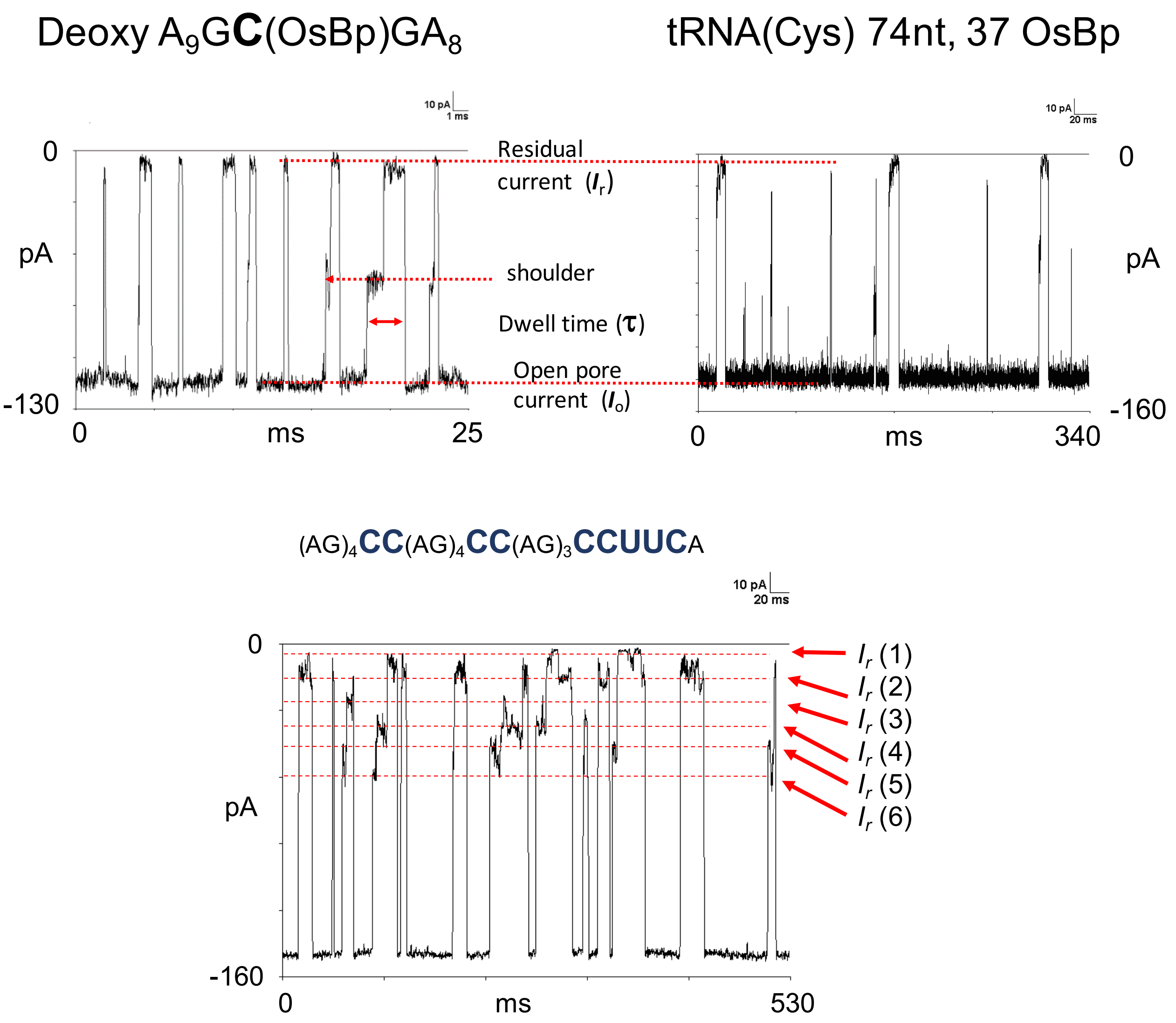


**Figure S18**. Concatenated *i-t* recordings from single translocations experiments via wt α-HL at 20ºC using the NanoPatch EBS instrument (see Experimental Section); electrolyte is 1M KCl, pH 7, 25mM TRIS.HCl buffer. **Top left:** deoxy $A_9G$**C**(OsBp)$GA_8$ at 120mV. **Top right:** osmylated tRNA (Cys) with 37 OsBp moieties at 160mV. **Bottom:** 32nt RNA oligo T8 (see sequence in Table 1, but with partial osmylation) at 160mV. Please note that the singly osmylated oligo exhibits a single level of residual current, whereas the translocations of the oligos with the multiple OsBp moieties exhibit more than one $I_r$ levels, consistent with rudimentary "sequencing" (see discussion in text). The multiple levels, maximum of 3 expected, are clearer to see with the 32nt RNA (bottom) due to scaling.

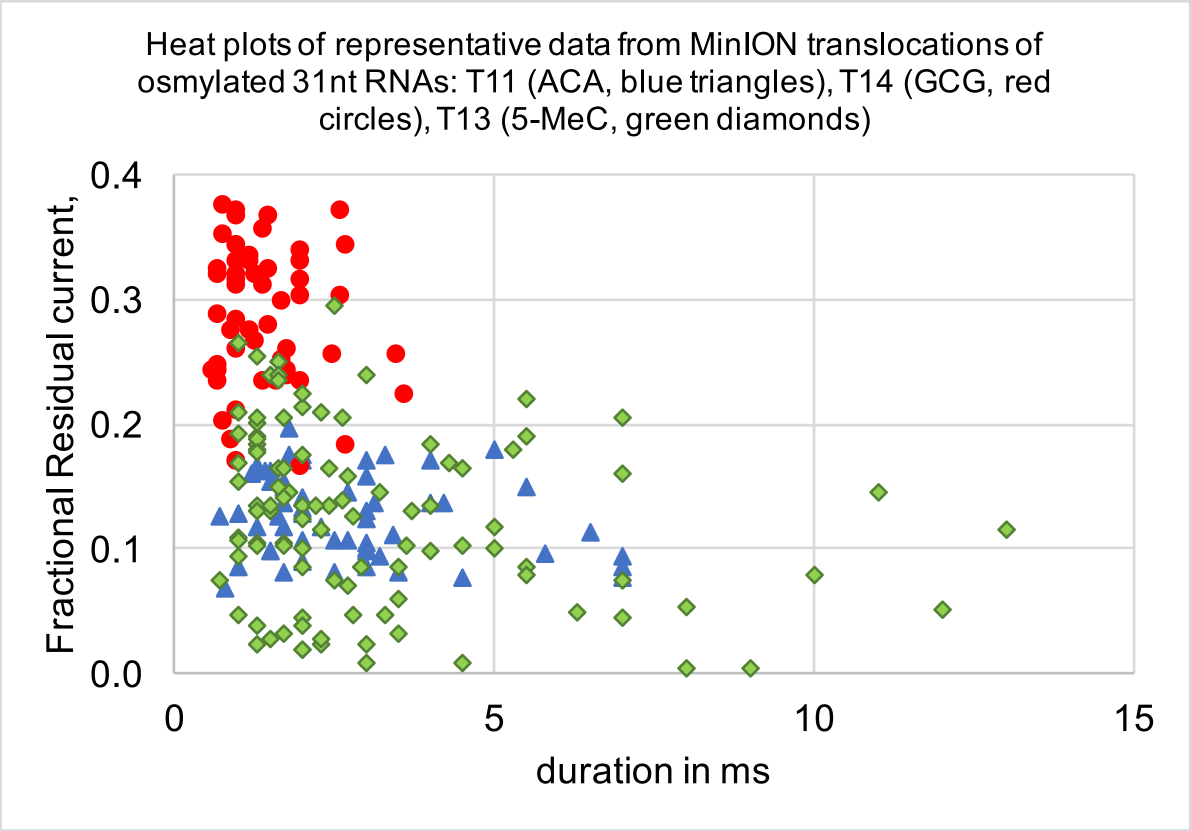


**Figure S19.** Heat plots of fractional residual ion current, $I_r/I_o$, as a function of duration, $\tau$ in ms, for three osmylated 31nt RNAs at -180mV. There is a definite trend of longer translocations with less residual ion current, $(I_r/I_o)_{max}$ (see Table 1 and text).
